# Air Pollution Particles Hijack Peroxidasin to Disrupt Immunosurveillance and Promote Lung Cancer

**DOI:** 10.1101/2021.12.17.473156

**Authors:** Zhenzhen Wang, Ziyu Zhai, Chunyu Chen, Xuejiao Tian, Zhen Xing, Panfei Xing, Yushun Yang, Junfeng Zhang, Chunming Wang, Lei Dong

**Author notes:** Corresponding Authors: L.D.,; C.M.W.,; J.F.Z.,.

## Abstract

Although fine particulate matter (FPM) in air pollutants and tobacco smoke is recognized as a strong carcinogen and global threat to public health, its biological mechanism for inducing lung cancer remains unclear. Here, by investigating FPM’s bioactivities in lung carcinoma mice models, we discover that these particles promote lung tumor progression by inducing aberrant thickening of tissue matrix and hampering migration of anti-tumor immunocytes. Upon inhalation into lung tissue, these FPM particles abundantly adsorb peroxidasin (PXDN) – an enzyme mediating type IV collagen (Col IV) crosslinking – onto their surface. The adsorbed PXDN exerts abnormally high activity to crosslink Col IV via increasing the formation of sulfilimine bonds at the NC1 domain, leading to an overly dense matrix in the lung tissue. This disordered structure decreases the mobility of cytotoxic CD8^+^ T lymphocytes into the lung and consequently impairs the local immune surveillance, enabling the flourish of nascent tumor cells. Meanwhile, inhibiting the activity of PXDN effectively abolishes the tumor-promoting effect of FPM, indicating the key impact of aberrant PXDN activity on tumorigenic process. In summary, our finding elucidates a new mechanism for FPM-induced lung tumorigenesis and identifies PXDN as a potential target for treatment or prevention of the FPM-relevant biological risks.

**Graphical abstract:** 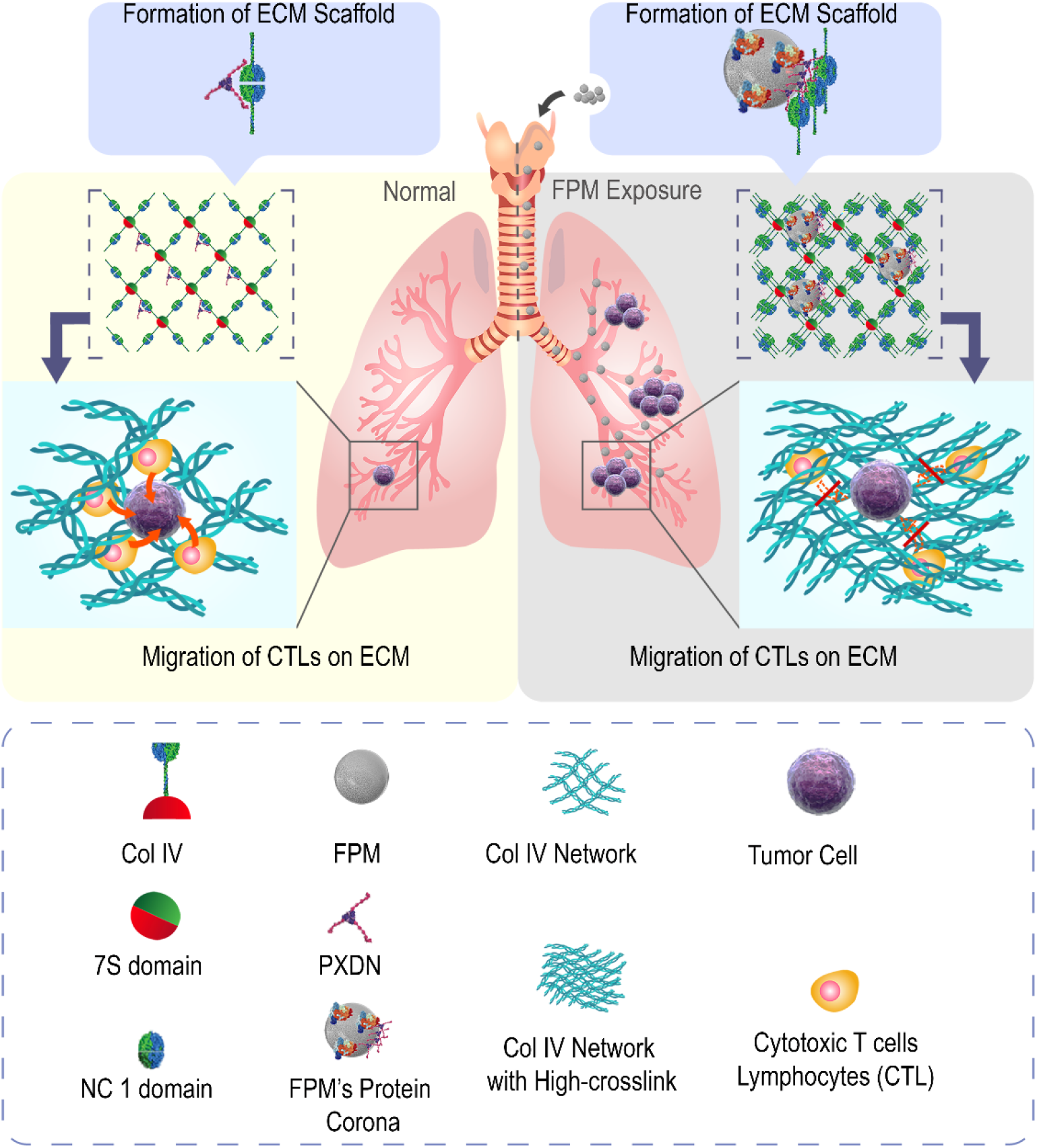

## Introduction

Inhalable fine particulate matter (FPM) with diameter less than 1 micron in air pollutants and tobacco smoke has been recognized as a Group 1 carcinogen and substantial threat to global health (36). About 10 μg/m^3^ increase in its concentration was correlated with 8% rise in lung cancer mortality (24). However, its carcinogenic mechanism remains unclear. During lung tumorigenesis, both the considerable growth of cancer cells per se and the accommodated supporting microenvironment are crucial (37). Earlier studies propose that the particles directly induce gene mutations and carcinogenesis (61), but recent investigations reveal that FPM does not directly promote (and even inhibit) the proliferation of lung cancer cells. These inconsistent findings suggest that FPM might have unidentified targets other than cancer cells in promoting tumorigenesis, such as immune cells that play key roles in tumor development (10). Under normal circumstances, the immune system rapidly detects and suppresses the tumor progression at the initial stage (28). Especially, the mostly ‘informed’ defender immunocyte - cytotoxic CD8^+^ T lymphocytes (CTLs) protect against potential cancer through efficient migration and cytotoxic contact with transformed or tumorigenic cells that have emerged in the lung interstitial space (15). Once this crucial immunosurveillance and defense process of CTLs were compromised, the tumorigenesis would be uncontrollable (30).

Under the chemotaxis of biochemical signals, the mobility of immunocyte depends on not only its intrinsic capacity but also the microstructure of interstitial extracellular matrix (ECM), that is, the way paved for immune cells (32). For the former, evidence about the direct effect of FPM on the immune cell’s migration capacity was validated. It’s estimated that tobacco smoke particulates (TSP) could impair the migration function of macrophage to mycobacteria and lead to increased susceptibility to tuberculosis in smokers (7). For the latter, clinical evidence links a dense collagen matrix surrounding the tumor with the restriction of T cells’ access (46; 53). Based on these reports and analysis, we speculated that FPM could disturb the migration and distribution of T cell in lung tissue, thus impairs CLTs’ immune defense capacity to cancerous cells, and consequently promotes tumor progression.

To test this hypothesis, we set out to study the effect of FPM inhalation on CTLs’ immune response and tumor development, by employing both transplantation (Lewis lung carcinoma, LLC) and transgenic (K-ras^G12D^p53^−/−^) mouse models of lung carcinoma (29). First, we validated that FPM promotes tumorigenesis by impairing CTLs’ migration towards cancerous cells. The defect was attributed to denser collagen structure induced by FPM on CTLs’ migration path, generating the physical isolation around tumor cells. More interestingly, we found that FPM exerts this effect by adsorbing peroxidasin (PXDN) – a crucial enzyme specifically mediating collagen crosslinking at NC1 domain – and increasing this enzyme’s activity to over-crosslink ECM and prevent CTLs migration, which eventually tolerates tumor progression.

## Results

### FPM Promotes Lung Cancer Development by Hampering CTLs’ Migration

To analyze the effect of fine particulate matter (FPM) on lung tumorigenesis, first, we collected and prepared particulate matter in air pollutants with diameter < 1 μm (PM1) from 7 locations in China and the tobacco smoke particle (TSP) respectively. Given that these particles displayed diverse morphology and physicochemical characteristic (**Supplementary Figure S1 and Supplementary Table S1**), which is consistent with the material of particles in other reports (31), we mixed PM1 with the equal proportion from each collection to eliminate the interference of sampling resources. Then, mice exposed to mixed PM1 (mixture) or TSP were analyzed on two cancer animal models: the syngeneic Lewis lung carcinoma (LLC) inoculation model (LLC-model) or the transgenic mouse model (Kras^G12D^p53^−/−^) as illustrated in **Supplementary Figure S2A**. Gross view (**Supplementary Figure S2B**) and histological hematoxylin and eosin (H&E) analysis (**Fig. 1A**) indicated that the FPM treatment markedly increased the tumors’ multiplicity and progression. As K-ras^G12D^p53^−/−^ mice could generate multifocal tumors corresponding to different grade of lung carcinoma (16; 47), histological grade of this model was further analyzed. Tumors in FPM-exposed lung tissue were mainly classified as grade II and the ones of grade III and IV were significantly higher, whereas the majority of tumors in the phosphate buffer saline (PBS) group were of grade I, showing FPM lead a more advanced lung tumorigenesis **(Fig. 1B)**. Statistical analysis suggested that the number of tumors in the FPM-treated group was significantly higher than that in the PBS group (about 3∼5 folds higher in LLC model and 2 folds higher in Kras^G12D^p53^−/−^ model) (**Fig. 1C**). Furthermore, the scenario was further validated by the corresponding tumor burden, based on the percentage of the area of tumor regions versus that of the total lung (about 7 folds more in LLC model and 10 folds more in Kras^G12D^p53^−/−^ model). Moreover, in both models, PM1 and TSP exposure significantly shortened the survival of mice (**Fig. 1D**). These results validated the correlation between FPM exposure and lung cancer development, in agreement with the epidemiological studies (5).

**Figure 1.**
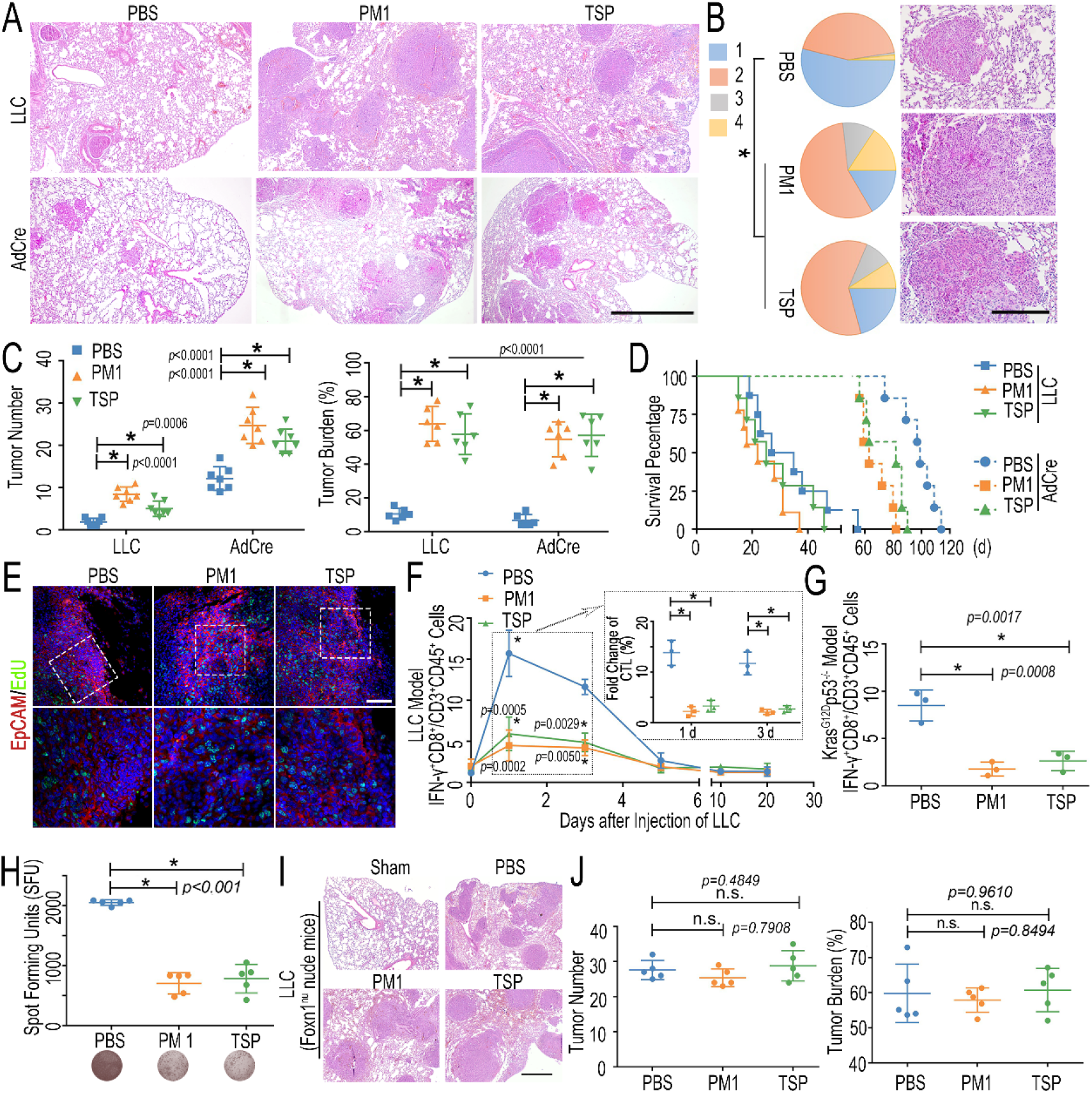
FPM accelerated lung tumorigenesis by inhibiting cytotoxic T cells lymphocytes (CTLs) infiltration. A. Representative hematoxylin and eosin (H&E) staining images of different model 20 days or 50 days after FPM-exposed mice were stimulated with LLC or Cre-inducible adenovirus (AdCre). Scale bar = 100 μm; B. Tumor stage (stages 1 to 4) of lungs in K-ras^G12D^p53^−/−^ model (left) and representative tumors H&E staining images with higher manification (right). *P* values are for comparisons of the percentage of stage 3 and 4 tumors in FPM-treated and control mice. n=5; C. Quantitative analysis about tumor number and tumor burden in lung tissue of different models treated as panel A. n=5; D. Survival analysis of mice exposed to FPM and subsequently stimulated with LLC or AdCre. n=7; E. Representative EdU staining images to analyze the proliferation rate of tumor site in lung tissue of LLC model 20 days after tumor initiation (DAPI, blue; EdU positive cells, green). Scale bar = 100 μm; F. The statistical analysis of CTLs (IFN-γ^+^CD8^+^/CD45^+^CD3^+^) in lung tissue based on flow cytometry at indicated day (0 d, 1 d, 3 d, 5 d, 10 d and 20 d) after intravenous injection of LLC. n=3. The inserted results in the dashed boxes indicated the fold change of CTLs in lung tissue 1 day and 3 days after the LLC stimulation, relative to that under the physiological condition; G. The statistical analysis of CTLs (IFN-γ^+^CD8^+^/CD45^+^CD3^+^) in lung tissue of K-ras^G12D^p53^−/−^ mice based on flow cytometry 4 weeks after after tumor initiation with the intratracheal injection of AdCre. n=3; H. IFN-γ enzyme-linked immunospot assay (ELISPOT) in the lung tissue of OT-1 TCR transgenic mice 1 day after OVA-LLC stimulation (upper) and represetative immunospot images (lower). n=5; I. Representative H&E staining images of lung tissue in *Foxn1^nu^* nude mice exposed to FPM 20 days after intravenous injection of LLC. Scale bar = 100 μm; J. Quantitative analysis about tumor number and tumor burden in lung tissue of *Foxn1^nu^* nude mice treated as panel I. n=5. Images are representative for three independent experiments. Results are shown as mean ± SD. \**p*<0.05 after ANOVA with Dunnett’s tests.

Next, we explored the reason for FPM promoting tumorigenesis. The conditions of the seeds and soil – the uncontrollably proliferative cancer cells and a tolerable immune microenvironment – are both crucial for tumor development (2). We analyzed the effect of FPM on the tumor cells and their congenial microenvironment, respectively. Interestingly, FPM hardly promoted the growth of tumor cells and even inhibited their proliferation at higher concentration (**Supplementary Figure S3).** EdU incorporation assay was further employed to determine the impact of FPM exposure on tumor cells’ proliferation *in vivo*. The result showed that the tumor site displayed similar replication capacity regardless of its size and advancement in these groups (**Fig. 1E and Supplementary Figure S4)**, casting the doubt of FPM’s direct promotion on tumor growth. These results inspired us to assess the effect of FPM on the immune microenvironment. Among these immunocytes related to immune surveillance, cytotoxic T lymphocytes (CTLs) as the most ‘‘informed’’ defender are critical for locally extinguishing the nascent tumor. The efficacy of these cells determines the fate of transformed cells – to death or flourish. Thus, we examined the change of CTLs’ response in different groups during tumor progression (**Fig. 1F and Supplementary Figure S5**). In the LLC model, CTLs in PBS group were efficiently recruited into lung tissue to defend LLC stimulation, increasing up to about 9-fold than that under the physiological condition at initial stage (1-3 days). Conversely, the lung tissue with FPM exposure displayed blunt and insufficient early immune defense, with slight CTLs infiltration, decreasing by more than 60% relative to that of PBS group, though there was no dramatic difference about CTLs accumulation among these groups at late stage (5-20 days). The immunofluorescence (IF) images also showed that FPM-exposed lung tissue was infiltrated with lower CTLs at initial stage (**Supplementary Figure S6**). These results indicated that the CTL’s early immune response might not be normal in FPM-exposed mice and be decisive for the lung tumorigenesis, which is consistent with the reports in transgenic autochthonous lung tumors (15). Then we detected the CTLs’ infiltration in Kras^G12D^p53^−/−^ model 4 weeks after tumor initiation, during which the immune response was reported to reach to the peak (15). The results displayed similarly insufficient CTLs’ defense in FPM-exposed group (**Fig. 1G and Supplementary Figure S7**).

To further determine whether CTLs’ reaction was specific to tumor cells, we further evaluated CD8^+^ T cell activation in an OT-1 TCR transgenic mouse model, in which the CD8^+^ T cells express a T cell receptor recognizing the SIINFEKL peptide of ovalbumin (OVA) (56). Upon the stimulation of OVA-expressing LLC (OVA-LLC) cells, the flow cytometry analysis of activated CTLs in lung tissue showed consistent tendency (**Supplementary Figure S8**). OVA-LLC specific immunity response in lung tissue was also tested by IFN-γ enzyme-linked immunospot assay (ELISPOT) (**Fig. 1H**). The result further demonstrated that the antigen specific activated CTLs was significantly impaired by FPM exposure, decreasing to about 25% of that in PBS group. Furthermore, to testify the indispensable role of CTLs on the tumor development, LLC cells were intravenously injected into FPM-exposed athymic *Foxn1*^nu^ nude mice with T cell deficiency. As expected, the difference of lung tumorigenesis in PBS and FPM-exposed groups was abolished (**Fig. 1 I, J and Supplementary Figure S9**), highlighting that the influences of FPM to tumor development were mediated by the CTLs. Additionally, without using tumor cells, we treated mice with a T cell chemokine – C-X-C motif chemokine ligand 10 (CXCL10) (23), or named as interferon-inducible protein-10 (IP-10) – intrabronchially for 2 h, as illustrated in **Supplementary Figure S10A**, and found that the proportion of CTLs in FPM-treated mice (about 8%) significantly decreased comparing with that of the PBS group (about 18%; **Supplementary Figure S10B**). This finding is consistent with the scenario observed in the tumor (LLC and Kras^G12D^p53^−/−^) model and strengthens the conclusion that FPM exposure delays the CTLs’ instantaneous defense response. The above results, taken together, indicate that FPM accelerates lung tumorigenesis *via* impairing CTLs infiltration into the lung tissue.

### FPM Hinders CTLs Migration by Crosslinking Type IV Collagen and Thickening Tissue Matrix

Next, we explored the reason for the impaired early response of CTLs under FPM exposure. CTLs distribution in the lung interstitial tissue depends on its migration ability (60), which is related to both the intrinsic activity of cells and the structure of the interstitial space formed by local extracellular matrix (ECM) on its migrating path (40). We analyzed which factor was mainly affected by FPM. First, T cells treated with FPM *in vitro* or the CTLs separated from FPM-exposed lung tissue were respectively analyzed. Integrin-1 (ITGB-1), C-X-C motif chemokine receptor 3 (CXCR 3) and Rho-associated kinase (ROCKi) (4; 41; 51), the biomarkers related to CTLs’ migration were detected with quantitative real-time polymerase chain reaction (qRT-PCR). These results showed that FPM stimulation had little effect on the migration potential of CTLs (**Supplementary Figure S11**). Second, we analyzed the change of the lung tissue structure after FPM exposure for 7 days. From SEM images and quantitative analysis about the pore size of interstitial matrix (**Fig. 2A and Supplementary Figure S12**), we noticed that the FPM exposure dramatically compressed the structure and crushed the interstitial space of the lung tissue. Further Massons trichrome staining indicated a higher density of collagen (**Fig. 2B**). These data implied that FPM inhaled into the lung tissue condensed the native framework of ECM, which could block the path of CTLs migrating to the tumor site.

**Figure 2.**
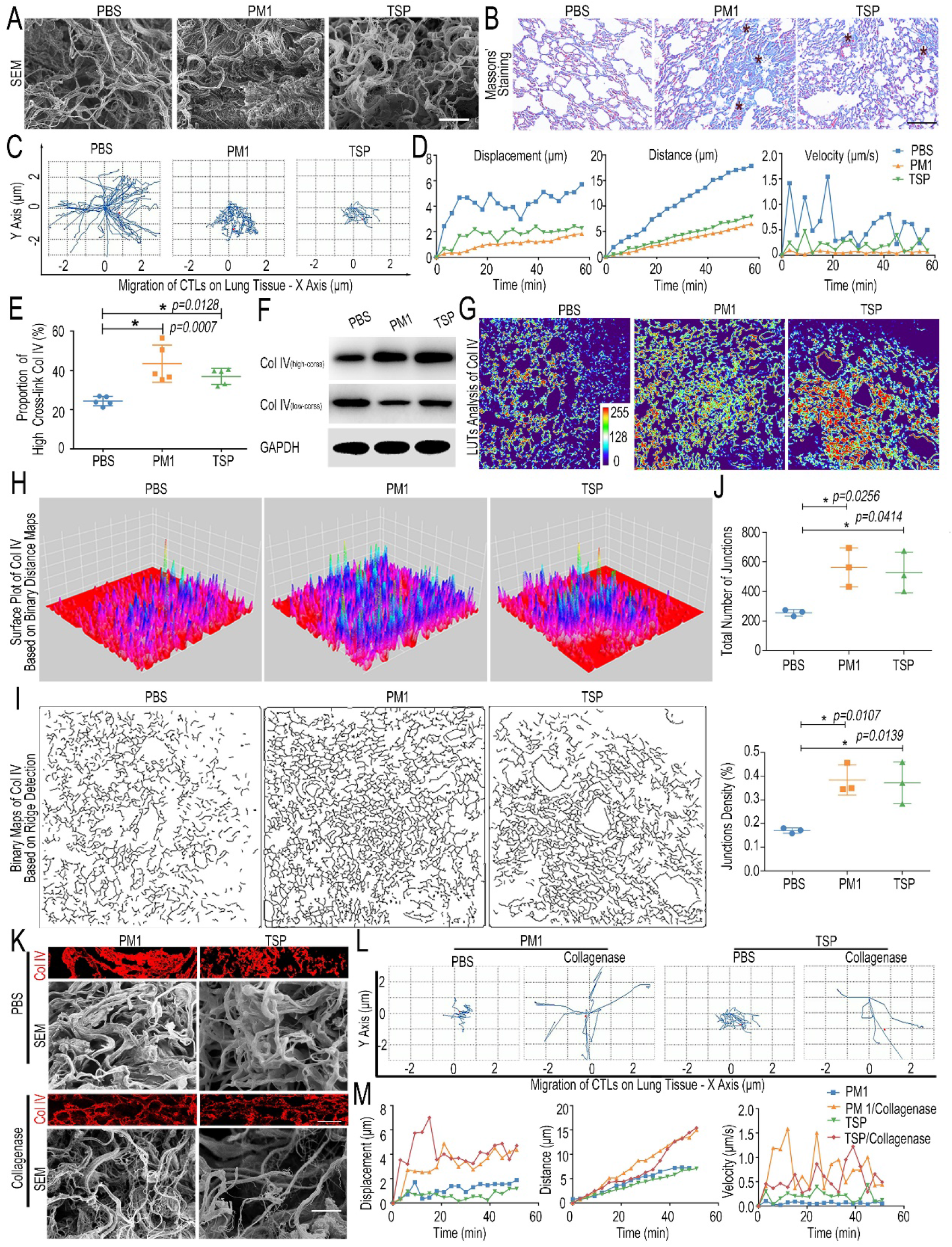
FPM impaired CTLs migration by increasing Col IV crosslinking in the lung tissue. A. Scanning electronic microscope (SEM) images of the interstitial matrix in the lung tissue exposed to FPM for 7 days. Scale bar = 100 μm; B. Representative Massons trichrome histological analysis of lung tissue exposed to FPM for 7 days. Images are representative of three independent experiments. Scale bar = 100 μm; C. Representative trajectory of CTLs’ migration in lung tissue slice of FPM-exposed mice or PBS group; D. The quantified analysis of migration distance, displacement and velocity of tracked CTLs *vs.* time (min) based on panel C; E. Proportion of high crosslink Col IV in lung tissue of mice exposed to FPM or PBS for 7 days, which was calculated by the ‘high-cross’ Col IV fragment divided by the sum of different fractions (‘low-cross’ ones and ‘high-cross’ ones). The content of each part was detected by ELISA. n=5; F. Western blotting analysis of ‘low-cross’ collagen and ‘high-cross’ collagen in lung tissue of mice exposed with FPM or PBS for 7 days; G-J. The in-depth analysis of representative Col IV immunofluorescence images of lung tissue in the mice exposed to FPM for 7 days through Image J: G. Look-up tables (LUTs) analysis of Col IV fluorescence intensity; H. Surface plot analyisis of Col IV distribution based on invert binary distance maps; I. Binary images of Col IV network generated by ridge detection plugin; J. Quantification analysis of junction number and denstiy in Col IV network based on panel I. n=3; K. Representative immunofluorescence images of Col IV and SEM images of FMP-exposed lung tissue treated with collagenase D (50 μg/ml). Scale bar = 10 μm; L. The trajectory of CTLs migrating in FPM-exposed lung tissue slice treated as panel K; M. Average migration distance, displacement and velocity of tracked CTLs *vs.* time (min) in lung tissue slice treated as in panel K. Images are representative for three independent experiments. Results are shown as mean ± SD. *p<0.05 after ANOVA with Dunnett’s tests.

Consequently, we investigated in greater detail the movement of CTLs in an *ex vivo* model. The migration of CTLs in the slice of lung tissue (native or FPM-exposed) was analyzed by dynamically visualizing the cells’ movement (**Supplementary Figure S13**). According to the outcomes from time-lapse sequential images and trajectory analysis of CTLs’ migration (**Fig. 2C and Supplementary Video S1-3**), in the lung tissue exposed to FPM, CTLs struggled to migrate, while those of PBS group displayed quick migration pattern. Statistical analysis further validated that comparing with the PBS group, the FPM-treated lung tissue severely hindered the migration of CTLs (**Fig. 2 D**), which were weakly motile and showed insufficient displacement, distance and velocity (11). Therefore, the change in the interstitial space, rather than attenuated migrating potential of CTLs per se, is responsible for the weakened infiltration of these cells in the FPM-treated lung tissue.

Then, we asked what caused the change of the lung structure after FPM exposure. Collagens, the main ECM components (21), especially 3 kinds of ones enriched in lung tissue, including the type I, III and IV ones (Col I, Col III and Col IV) were focused on (33). First, enzyme-linked immunosorbent assay (ELISA) was performed after different fragments of collagens were respectively harvested and divided into 2 categories by a reported protocol (44), that is, ‘low-crosslinked’ ones (low-cross) – containing freshly secreted collagens, procollagens and moderately crosslinked collagens – and the other remainder ‘high-crosslinked’ ones (high-cross) (**Supplementary Figure S14**). The results showed both PM1 and TSP exposure significantly elevated the high-crosslink proportion of Col IV, 2-fold higher than that in PBS group, based on the separate examination of low-cross and high-cross ones (**Fig. 2 E**). Next, the relative quantification of high-crosslinked collagens compared with low-crosslinked ones based on western blotting analysis showed consistent changes (**Fig. 2 F and Supplementary Figure S15**). Besides, the IF images indicated that FPM exposure induced Col IV in the lung tissue to generate enhanced crosslink and denser distribution, leading the collagen network with more junction site and higher junction density (**Fig. 2 G-J and Supplementary Figure S16**). However, the other two types of collagens (Col I and Col III) showed no obvious change (**Supplementary Figure S17**), demonstrating an increased Col IV crosslinking accounted for the change in the lung ECM structure. Furthermore, based on the related integrated optical density (IOD) of Col IV in lung tissue slices and related CTLs’ migration index (migration distance, displacement and velocity) of different groups in **Figure 2D**, we performed the Pearson’s correlation analysis and found an inverse relationship between Col IV density and CTLs’ migration potential (**Supplementary Figure S18**). Furthermore, we pre-treated the FPM-exposed lung tissue with collagenase D to reduce the Col IV crosslink and alleviate Col IV density (**Fig. 2 K**). The trajectory images and related quantification analysis showed CTLs’ migration was effectively reversed (**Fig. 2L, M and Supplementary Video S4-7**), further validating the crucial role of Col IV crosslink on the CTLs’ movement. These data together suggested that FPM exposure blocked CTLs migration and trapped these cells mainly through increasing Col IV crosslinking and consequently generating a denser ECM in the lung tissue, which might isolate the tumor cells from the CTLs’ attack (**Supplementary Figure S19**).

### FPM Increases Col IV Crosslinking through Promoting Sulfilimine Bond Formation

We then investigated why FPM exposure led to increased Col IV crosslinking (21). According to recent discovery, protein adsorbed onto the nanoparticles surface would endow them with new activities (39; 57). As the median size of the both PM1 and TSP is about 100-200 nm, it is possible that the collagen-crosslinking activity of FPM is derived from the proteins adsorbed onto their surface from lung tissue. To elucidate this, we separately incubated FPM in PBS or lung homogenate (LH) to simulate the scenario of FPM per se (FPM group) or the complex of FPM and its surface proteins (LH-FPM group, including LH-PM1 and LH-TSP). Then, according to an established experimental model with a slight modification (9), soluble Col IV was generated by stimulating mouse bone marrow fibroblasts M2-10B4 cells, which highly express Col IV, with the inhibitor of collagen crosslink (**Supplementary Figure S20**). Next, the effect of FPM on the crosslink of soluble Col IV in the cellular system and acellular system was respectively analyzed shown as **Fig. 3A**. For the cellular system, Col IV immunostaining result of M2-10B4 cells showed FPM itself could not induce the crosslinking of soluble Col IV (**Fig. 3B**). Relatively, the LH-FPM initiated the crosslinking and reinforced the network to a greater extent than that induced by LH per se, which could be validated by intensive crosslink intensity and a denser Col IV distribution, and collagen network with more junction site and higher junction density (**Fig. 3C-F**). Meanwhile, for the acellular experiment, the cell lysate of M2-10B4 enriched soluble Col IV was incubated with FPM or LH-FPM mixture. Western blotting result showed a similar scenario – the naked FPM had little devotion to Col IV crosslink, but the LH-FPM dramatically enhanced the high-crosslinked Col IV fragment (**Fig. 3G and Supplementary Figure S21**).

**Figure 3.**
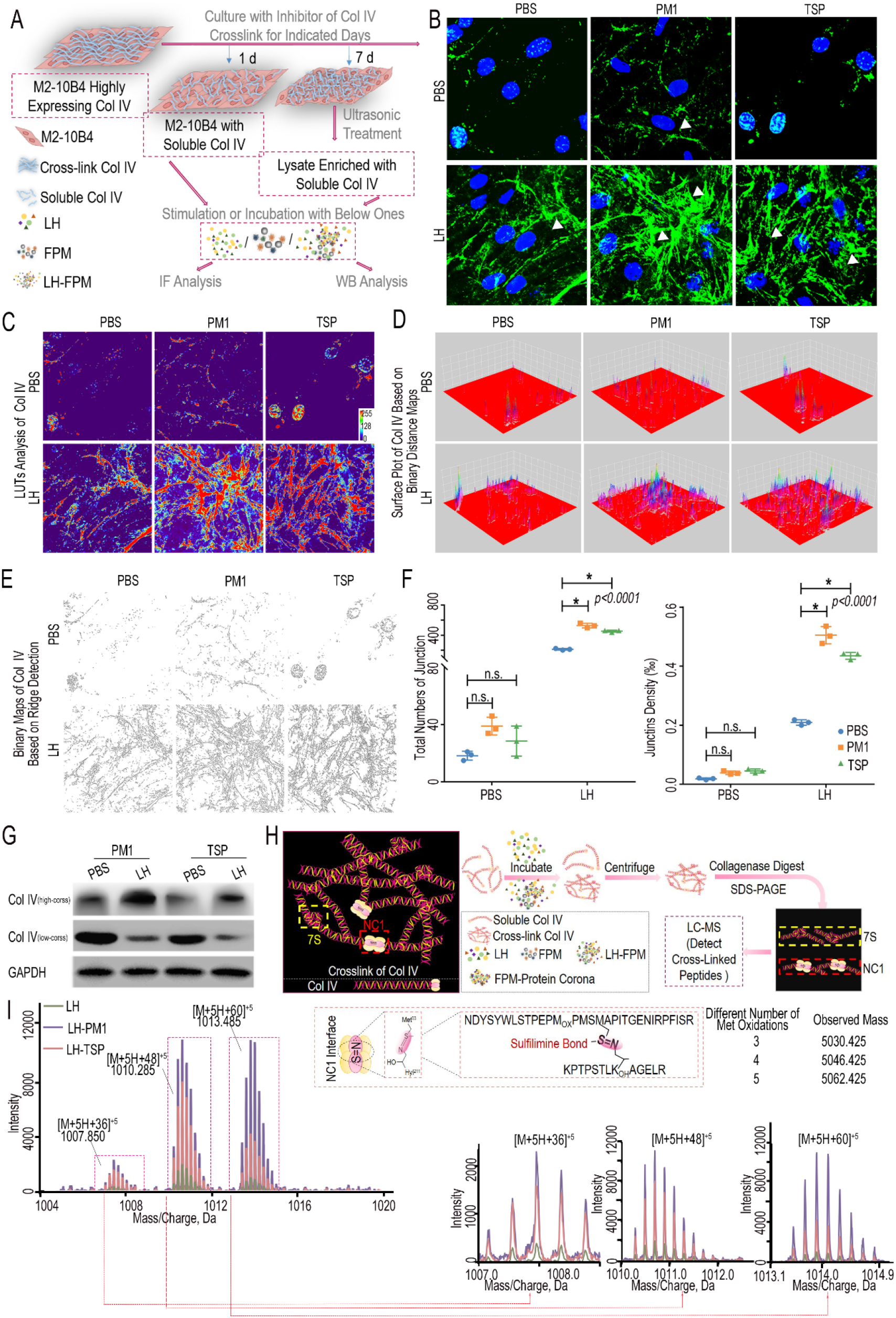
FPM increased Col IV crosslink *via* promoting sulfilimine bond formation at the NC1 domain. A. Schematic representation of the procedures to generate soluble Col IV and analyze the effect of FPM on its crosslink. Briefly, the M2-10B4 cells highly expressing Col IV were treated with crosslink inhibitor for 1 day or 7 days and then treated with FPM or the mixture of LH-FPM, stimulating the scenario of FPM *per se* or its interface with LH, to initiate the crosslink. The crosslink level of Col IV was analyzed with diversity methods; B. Representative immunofluorescence capture of Col IV in M2-10B4 cells stimulated with FPM or LH-FPM for 24 h after pretreated with crosslink inhibitor for 1 day. Scale bar = 20 μm. Images are representative of three independent experiments; C-F. The in-depth analysis of Col IV immunofluorescence images in panel B through Image J: C. Look-up tables (LUTs) analysis of Col IV fluorescence intensity; D. Surface plot analysis of Col IV distribution based on invert binary distance maps; E. Binary images of Col IV network generated by ridge detection plugin; F. Quantification analysis of junction number and denstiy in Col IV network based on panel E; G. Western blotting of ‘low-cross’ and ‘high-cross’ Col IV fraction in M2-10B4 cells lysate enriched with soluble collagen after their treatment with FPM or LH-FPM; H. Schematic diagram of separating fragments containing the NC1 domain crosslink site in Col IV. The general crosslink network generated by Col IV was displayed on the left, with the important crosslink sites (7S domain and NC1 domain) respectively labelled in the yellow and red dotted box; I. High-resolution mass spectrum depicting tryptic peptides containing sulfilimine bond (-S=N-), with magnified spectrum displayed on the bottom. The formation of -S=N- and the known peptide sequence with different oxidation containing the sulfilimine bond were shown on the upper right.

These data raised the question that how LH-FPM increased Col IV crosslinking. During crosslinking, the triple-helical protomer of Col IV, as the building block, form network through two key types of crosslinking sites (12): NC1 domains including sulfilimine bond (- S=N-) formed at the *C*-terminal (54) and 7S tetramers including aldehyde group formed at the *N*-terminal (45) (**Fig. 3H**). To distinguish which one is mainly disturbed by the FPM, we detected their changes under FPM stimulation respectively: 1) For the NC1 domain, sulfilimine bond (-S=N-) is formed by two juxtaposed Col IV protomers at residues methionine 93 (Met93) and hydroxylysine 211 (Hyl211) (**Supplementary Figure S22**) (9). Based on indicated theoretical mass of crosslinked tryptic peptides containing -S=N- in NC1 domain, we performed high-resolution liquid chromatograph-mass spectrometer (LC-MS) analysis to differentiate these peptides. For the NC1 domain separated from the crosslinked Col IV as illustrated in **Fig. 3H**, we found significantly more sulfilimine-containing peptides in LH-FPM-treated soluble Col IV than that in LH group according to the total ion chromatography (TIC) diagram (**Fig. 3I**). 2) For the 7S domain crosslinking site, it was derived from the oxidation of one lysine residue in the N-terminal to the aldehyde (3). The generated allysine would subsequently undergo a series of condensation reactions with other amino acids, mainly the other lysine or lysines on neighboring C-terminus, forming methylenimine bond (-C=N-), pyridine or others to stabilize crosslink (**Supplementary Figure S23A**). During the process, the detection of primary product allysine could reflect the level of 7S domain crosslink. With the reported specific and efficient probes to allysine (55), the allysine yielded during the crosslinking of the soluble Col IV incubated with LH or LH-FPM were respectively analyzed. The result showed a slight difference, indicating the 7S domain would not be interfered by FPM (**Supplementary Figure S23B**). Summarily, FPM gained a catalyzing activity from the proteins adsorbed from the tissue, which mediated the cross-linking through forming excessive -S=N-bonds among Col IV molecules.

### Phase Transition of PXDN on FPM Surface Increases Its Activity for Col IV Crosslinking

Although the above findings demonstrated that LH-FPM increased sulfilimine bond formation to enhance Col IV crosslinking, it remained unclear how FPM gain the activity from LH to mediate this biochemical process *in vivo*. To elaborately dissect this process, following a standard procedure (**Supplementary Figure S24**) (13), we separated the biomacromolecules from LH-FPM, the majority of which are proteins and also known as ‘protein corona’ (39). Unexpected corona formation can trigger serious pathological reactions (50; 57). Thus we speculated whether FPM could recruit certain proteins related to Col IV crosslink into its corona, thereby enriching and empowering this protein – to influence the crosslinking of collagen IV.

Given that the sulfilimine bond is uniquely catalyzed by peroxidasin (PXDN) enzyme in animal tissue (9; 58), we focused and detected the PXDN in FPM’s protein corona. The LC-MS result showed that PXDN was listed in the component profile of protein corona on both PM1 and TSP (**Supplementary Table S2**), reflecting the interaction of PXDN and FPM. We further analyzed PXDN adsorbed on FPM and its time evolution with Western blotting (57). The data showed that PXDN was not only enriched on the FPM (**Fig. 4A)**, but also stably tethered to FPM as the incubation time increased (**Fig. 4B)**, underlying it might affect FPM’s biological behavior durably. To further assess the adsorption of FPM to PXDN, we injected rhodamine fluorescence-labelled particles (R-FPM) *via* trachea into the lung tissue and detected PXDN therein. The IF co-localization of FPM and PXDN *in vivo* was clearly presented, confirming the recruitment of this enzyme to FPM (**Fig. 4C**), which could be further validated by the dramatically similar distribution of PXDN and FPM on M2-10B4 cells (**Figure 4D**). Taken together, these data indicate that FPM enriches and stabilizes PXDN in its surface corona.

**Figure 4.**
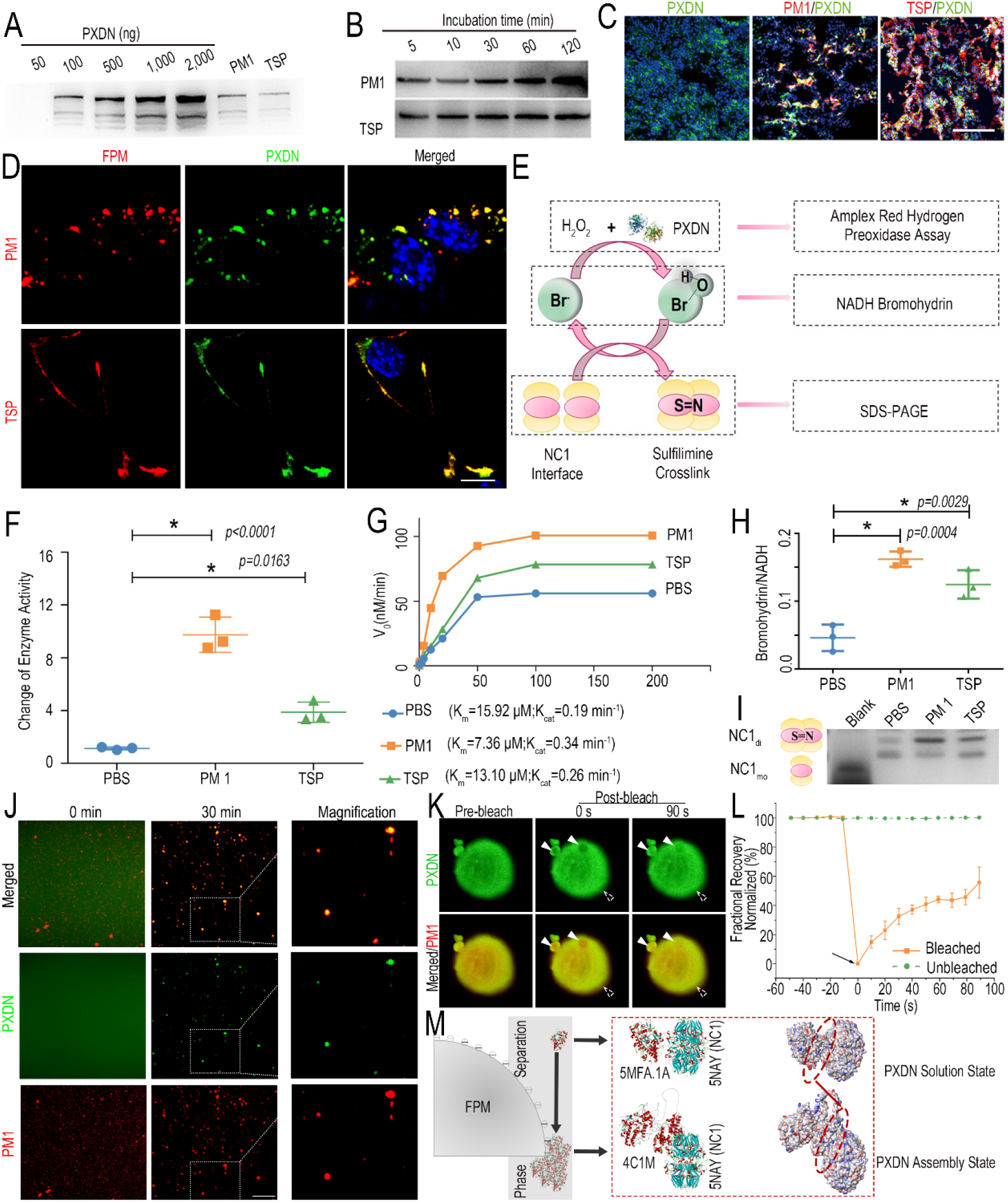
FPM increased PXDN activity by triggering the enzyme’s phase transition. A. Quantitative western blotting analysis of PXDN harvested from FPM corona formed in LH after incubation for 2 h, with a serial content of recombinant PXDN protein as the standard control; B. Western blotting analysis of PXDN at indicated time points to identify its time evolution in FPM’s protein corona; C. Representative confocal microscopic photographs showing the co-localization of rhodamine-labelled FPM (shown in red) and PXDN in lung tissue. PXDN was indicated as green. Scale bar = 100 μm); D. Representative fluorescent photographs of M2-10B4 cells treated with rhodamine-labelled PXDN and FITC-labelled FPM for 1 h. Rhodamine-labelled PXDN was shown in red and FITC-FPM in green. Scale bar = 10 μm; E. Schematic representation of the procedure about the formation of sulfilimine bond catalyzed by PXDN. Aimed at the substrate H_2_O_2_, the intermediates HOBr and the final NC1 domain with sulfilimine bond, different experimental analyses were respectively performed; F and G. Fold change of enzyme activity (F) and enzyme kinetics (G) of PXDN stimulated with FPM, determined by Amplex Red Hydrogen Peroxidase Assay Kit; H. Ration of NADH bromohydrin relative to NADH based their intensity of peaks detected by LC-MS. The analysis was performed after PXDN was incubated with FPM for 30 min and then catalyzed in the presence of 100 μM H_2_O_2_ and 200 μM NaBr at 37°C for 30 min; I. SDS-PAGE and Coomassie staining of NC1 domain 4 h after they were incubated with PXDN, following the latter’s incubation with PBS or FPM for 30 min. The crosslinked dimeric (NC1_di_) and un-crosslinked monomeric subunits (NC1_mo_) respectively labelled. Images are representative of three independent experiments; J. The confocal microscopy of FITC-labelled PXDN incubated with rhodamine-labelled FPM in LH for the indicated time (0 min and 30 min). Shown at the right were images with higher magnification for the assemblies of PXDN’s liquid-like droplets on the FPM at 30 min. Scale bar = 5 μm; K. Representative images from FRAP experiments showing the dynamic and reversible characteristic of PXDN-droplets. The rhodamine-labelled PXDN was shown in red and FITC-labelled FPM in green. The bleached region of interest (ROI) was indicated with white triangles and the unbleached control ROI was labelled with dotted white ones; L. Quantification of fluorescence recovery percentage in the ROI regions of PXDN’s liquid-like droplets. The black arrow indicates the initiation of laser bleach treatment; M. Interactive docking model on the effect of PXDN’s phase separation on its enzymatic performance at the catalytic interface of NC1 domain. The structure of PXDN solution state (PDB ID: 5MFA.1) and its assembly (PDB ID: 4C1M; created through homology modeling) were respectively displayed as the lateral stereo view of transparent chain model (left) and Dand swiss model (right). The interface site of contact between NC1 domain (PDB ID: 5NAY) and PXDN was labelled with the red dotted ellipses. n=3. Results are shown as mean ± SD. \**p*<0.05 after ANOVA with Dunnett’s tests.

The adsorption of FPM might disturb the activity of proteins, especially for the enzyme therein. Thus, we examined the activity of FPM-recruited PXDN shown as illustrated in **Fig. 4E**. Aimed at the substrate H_2_O_2_, the intermediates hypohalous acids and the final NC1 domain with sulfilimine bond, different experimental analyses were respectively performed (9). First, peroxidase activity of PXDN incubated with FPM was measured through Amplex Red molecular probes (6). The relative fluorescence intensity showed that FPM incubation raised the enzymatic activity of PXDN up to 5-10 folds (**Fig. 4F**). Next, PXDN’s enzyme kinetic behaviors were investigated according to the Michaelis–Menten model. Based on the generated Line weaver–Burk representative plot, the Michaelis constant (Km) and turnover number (Kcat) were calculated, which respectively reflects the enzyme-substrate binding efficiency and the enzymatic efficiency. The result showed that PXDN incubated with FPM revealed significantly decreased Km (PM1: 7.36 μM; TSP: 13.10 μM *versus* PBS: 15.92 μM) and increased catalytic efficiency about 1.4-to-1.8-fold than the enzyme per se (**Fig. 4G**). Also, the secondary product hypohalous acids (9; 38), for instance HOBr and HOCl, generated by PXDN from bromide and chloride was respectively analyzed. For HOBr, bromohydrin formed by the bromination of NADH was measured based on its close relation with HOBr production as reported in the literature (6; 49). The LC-MS detection showed that the ratio of bromohydrin to NADH in the group of PXDN incubated with FPM was 3-5-fold higher than that of the enzyme per se group, indicating more HOBr production (**Fig. 4H and Supplementary Figure S25**). Besides, we measured HOCl in consideration of the vast excess of Cl^−^ over Br^−^ in most animals (59), although PXDN uses Br^−^ to catalyze formation of sulfilimine crosslinks with greater efficiency (38). With the detection of a hypohalous acid-detecting fluorescent probe (62), we found HOCl produced by the FPM-incubated PXDN was about 2-to-4-fold higher than that of PBS group (**Supplementary Figure S26**). Furthermore, to analyze the effect of FPM’s adsorption on the catalytic product, after the commercial non-crosslinking NC1 domain (NC1_mo_) was reacted with FPM-incubated PXDN, the generated crosslinked dimeric with sulfilimine bond (NC1_di_) therein were detected by SDS-PAGE (9; 19). The result suggested that FPM incubation significantly enhanced PXDN’s enzymatic performance, with a higher yield of crosslinked NC1 dimeric subunits (**Fig. 4I**). Taken together, these results suggest that the pro-crosslink potential of FPM attributed to the aberrant enzyme activity of PXDN adsorbed on its surface.

Next, we gave an insight into the mechanism of PXDN’s tampered catalysis. Recent emerging evidences suggest that phase transition (or separation) is a common way to regulate proteins’ activity (27). Many physiochemical factors interfere or fluctuation could initiate proteins’ phase separation. And we doubt whether the FPM’s adsorption could induce the PXDN’s phase transition, thus disturbing the latter’s enzymatic activity. Taking PM1 as an example, with a series of microscopic observation, we found the formation of PXDN’s liquid-like droplets in LH after its incubation with PM1 with both confocal fluorescence and phase contrast microscope (**Fig. 4J, Supplementary Figure S27A**). However, for the PXDN alone in LH under the same processing time and imaging parameters, no assemblies are observed. Moreover, the profiles of fluorescence distribution further indicated the phase-separating PXDN’s accumulation was initialized on the PM1, based on their evident colocalization (**Supplementary Figure S27B**). Besides, to characterize the dynamic nature of the droplets, we performed fluorescence recovery after photobleaching (FRAP) experiments. FRAP studies revealed that fluorescence recovery of the bleached region of PXDN droplets could be partially recovered in minutes after photobleaching (**Fig. 4K and L**). The reversible characteristic observed for PXDN droplets on PM1 further validated that PXDN underwent phase separation. More interestingly, the droplet-like accumulation of PXDN on FPM could be also observed in M2-10B4 cells (**Figure 4D**). Overall, these results indicated that the increased activity for PXDN’s crosslinking Col IV was triggered by its phase transition on FPM surface.

Furthermore, we theoretically inferred the relationship of PXDN’s phase separation with its enzymatic performance. First, we focused on the low complexity domains in PXND sequence, which could drive phase transition and be predicted by intrinsically disordered regions (IDR; **Supplementary Figure S28**). The analysis indicated PXDN’s phase separation might occur at sequence 200-400 aa, which shows higher IDR score (1). More importantly, it’s away from PXDN’s activity center (800-1200 aa) according to the spatial structure in SWISS-MODEL, giving us a hint that PXDN’s phase transition on FPM surface might not lead to deformation of PXDN’s catalytic center and loss of its function. Next, protein-protein interactive docking simulation were performed at the interface of PXDN with its substrate NC1 domain through ZDock protocol (**Figure 4M**). As a homotridmeric multidomain enzyme, PXDN exists as trimerization in solution, and three monomers of PXDN are linked by disulfide bonds at the indicated flexible linker region in the residual non-catalytic domains (6). According to the reported modelled structure of PXDN (42), its trimeric peroxidase domain displayed a triangular arrangement. However, oversized trimerization of PXDN might not get in contact thoroughly with its substrate (34). So we chose the PXDN monomer with the exposed enzymatic surface contacting with NC1 domain to simulate its function at solution state. The computational result indicated that once PXDN triggered phase separation, which transferred from the solution state to the aggregation one (the putative assembly structure created through homology modelling) (8), the interactive area at the NC1 interface would be notably increased, thus facilitating the enzymatic catalysis.

### Inhibiting PXDN Ameliorates FPM-Induced Tumorigenesis

Based on the above findings, we speculated that inhibiting PXDN could abolish ECM change and recover CTLs migration in the lung. To testify our hypothesis, the effect of the plasmids capable of ectopically expressing PXDN specific short hairpin RNA (shPXDN) was detected. After validating its interference efficiency (**Supplementary Figure S29**), shPXDN mixed in the *in vivo*-jetPEI gene transfer regent was delivered into murine lung tissue through trachea injection, as illustrated in **Supplementary Figure S30**. The assessment of Col IV with different fraction (‘low-cross’ ones and ‘high-cross’ ones) measured by ELISA revealed the shPXDN diminished crosslinked level of Col IV (**Fig. 5A**). Massons trichrome staining images and SEM observation also confirmed that the shPXDN effectively decreased collagen density and expanded interstitial space (**Supplementary Figure S31**). More importantly, the ameliorative Col IV network induced by shPXDN further recovered the migration and accumulation of CTLs in lung tissue 1 day after the LLC stimulation, revealed as the CTLs’ migration trajectory images and the immunofluorescent staining (**Fig. 5B and C**). Flow cytometry analysis further demonstrated that shPXDN efficiently reversed the CTLs’ infiltration in the FPM-exposed lung tissue (**Fig. 5D and Supplementary Figure S32**).

**Figure 5.**
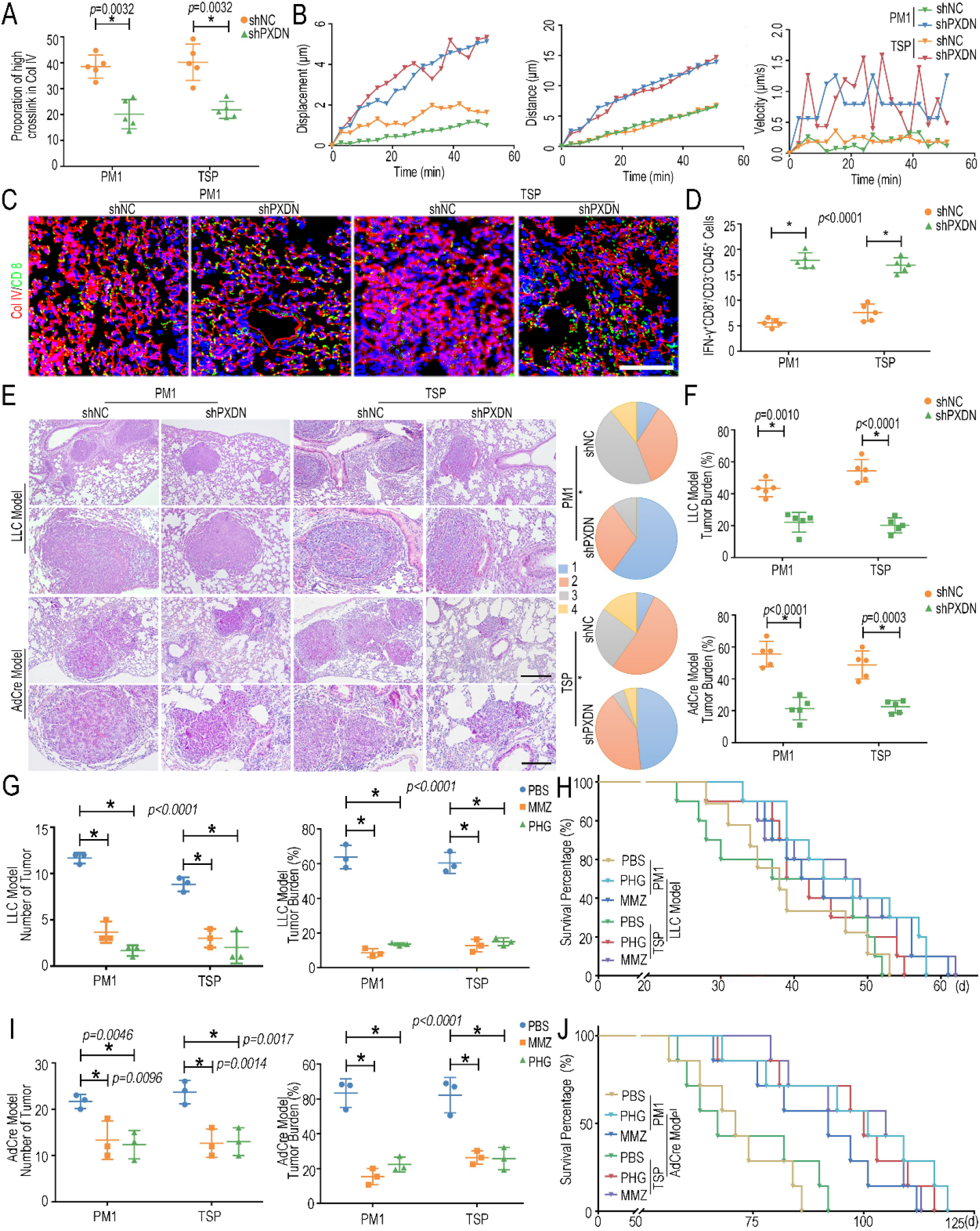
PXDN inhibitor alleviated FPM-induced lung tumorigenesis. A. Proportion of high crosslink Col IV in lung tissue of FPM-exposed mice pretreated with PXDN specific short hairpin RNA (PXDN shRNA), which was calculated by the ‘high-cross’ Col IV fragment divided by the sum of different fractions (‘low-cross’ ones and ‘high-cross’ ones) based on ELISA; B. Migration distance, displacement and velocity *vs.* time (min) of tracked CTLs in lung tissue slice of FPM-exposed mice administrated with PXDN shRNA; C. Representative immunofluorescence images of CTLs’ infiltration into the FPM-exposed lung tissue pretreated with PXDN shRNA 1 day after intravenous injection of LLC. Scale bar = 100 μm; D. The statistical analysis of CTLs (IFN-γ^+^CD8^+^/CD45^+^CD3^+^) based on flow cytometry in lung tissue of mice treated as in panel C. n=5; E. Representative H&E staining images of lung tissue (the lower ones: capture with higher magnification) yielded from LLC model and Kras^G12D^/p53^−/−^ model after mice pretreated with PXDN shRNA. The scale bar = 200 μm (upper) and 100 μm (lower). Tumor stage (stages 1 to 4) in lungs of K-ras^G12D^p53^−/−^ mice was shown in the right. *p* values are for comparisons of the percentage of stage 3 and 4 tumors in different group. n=5; F. Statistical analysis about tumor burden of mice in LLC model and Kras^G12D^/p53^−/−^ model administrated with PXDN shRNA. n=5; G-J. Statistical analysis about number and burden of tumors and survival curve of mice in LLC model (G and H) and Kras^G12D^/p53^−/−^ model (I and J) administrated with methimazole (MMZ) or phloroglucinol (PHG). Images are representative of three independent experiments. n=3. Results are shown as mean ± SD. *p<0.05 after ANOVA with Dunnett’s tests.

These results encouraged us to assess the effect of shPXDN in suppressing the tumor progress induced by FPM, as illustrated in **Supplementary Figure S33**. Encouragingly, our data showed that the shPXDN significantly suppressed tumor growth, diminished tumor grade, and reduced tumor number and burden (**Fig. 5E, F and Supplementary Figure S34**) in both K-ras^G12D^p53^−/−^ transgenic model and LLC-model. These results suggested the feasibility of inhibiting PXDN as a potential therapeutic target for FPM-associated lung cancer. Moreover, the efficacy of small molecule PXDN inhibitors, including methimazole (MMZ) and PHG (9), were similarly studied with optimal therapeutic doses (**Supplementary Figure S35**). To be satisfactory, both MMZ and PHG also effectively suppressed lung tumorigenesis (**Fig. 5G-J and Supplementary Figure S36**), further expanding the strategy for lung cancer treatment.

## Discussion

Although inhalable particles from air pollutants and tobacco smoke have clearly been identified as a potent carcinogen to humans, its pathological mechanism remains unclear, which hampers the design of therapeutic approaches. Existing findings, though supporting that fine particulate matter (FPM) induces lung cancer, provide insufficient and inconsistent explanations for the underlying mechanism. In the present study, we have discovered an unexpected mechanism that, as shown in the **Fig. 6**, apart from directly targeting tumor cells, the inhaled FPM changes the formation of lung tissue matrix to prevent the infiltration of T lymphocytes and their anti-tumor immunosurveillance, which consequently accelerates lung tumorigenesis.

**Figure 6.**
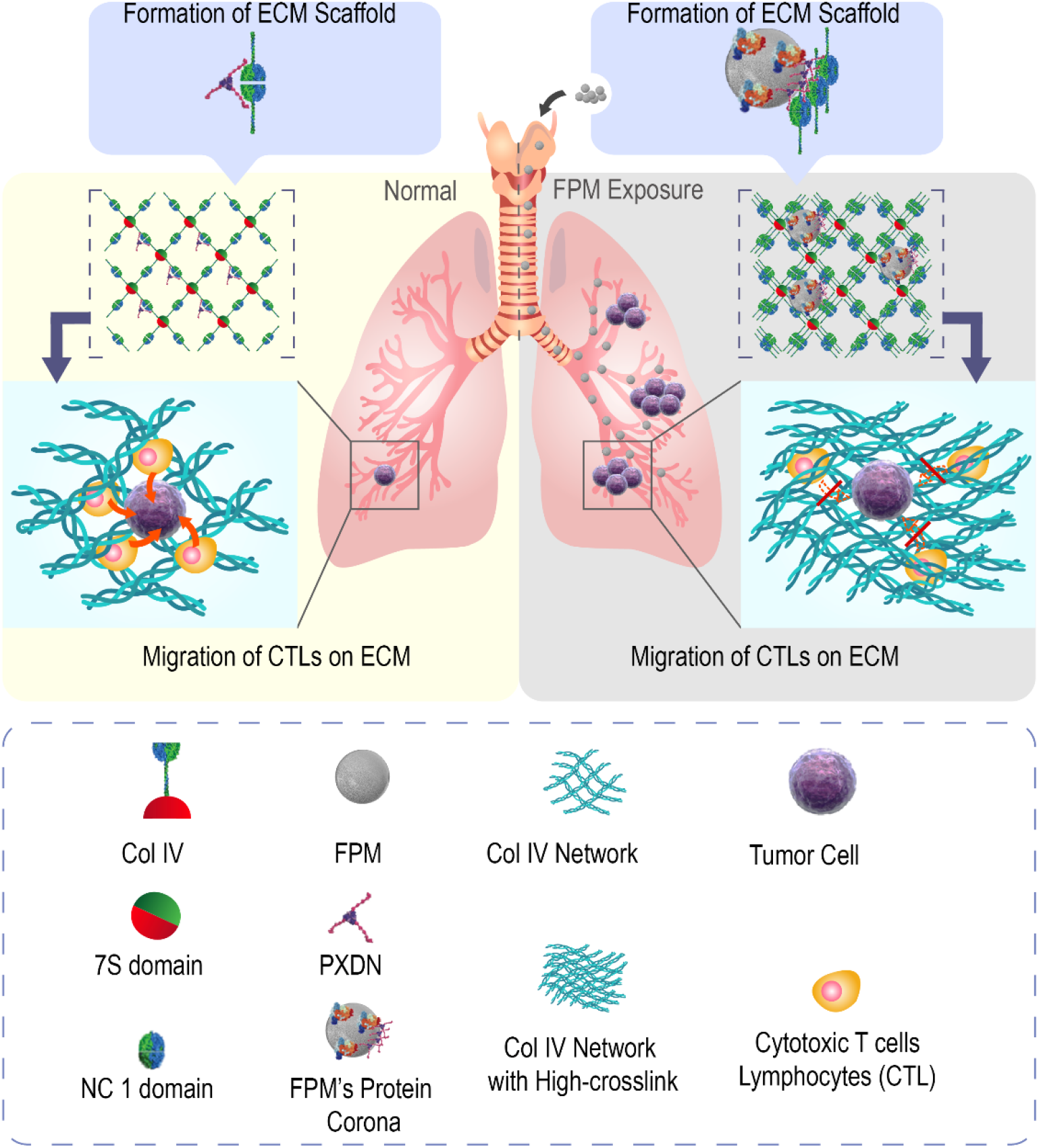
Schematic diagram of the mechanism underlying how FPM promotes lung tumorigenesis. The adsorption of FPM triggers the phase separation of PXDN to generate an aberrant catalytic activity and induce disordered crosslink of Col IV. Reinforced Col IV network impaired CTLs migration and local immunosurveillance, which considerably increases tumorigenesis in lung tissue.

Our study was inspired by an unanswered, fundamental question – which component in the lung tissue is first affected by FPM? Most previous studies suggest that FPM directly act on cells, by inducing gene mutation and increasing malignant ‘seed cells’ in the affected lung (48). They further suggest that the particles carry chemical reagents, such as acrolein, nicotine, oxidants and reactive nitrogen moieties (26), into the lung to exert such effects. However, the respiratory tract is exposed to various environmental substances (e.g. O_2_ or O_3_) that are much more powerful in damaging DNA, more abundant in the air, and more soluble in the tissue than the above chemicals. It requires further explanation how these assumed mutagens, delivered in a trace amount by the particles into the lung tissue, could significantly increase the lung cancer incidence. Moreover, FPM from different locations carries distinct types of those carcinogens exiting in the local environment, but these highly various particles exert the same carcinogenic effect (31), which is unexplainable. Additionally, *in vitro*, FPM is considered more capable of entering the cell than larger particles (22) (such as the PM10, that is, the particulate matter with the diameter of 1-10 μm); but *in vivo*, particles are difficult to directly contact with cells, due to the barrier effects from sticky body fluids and gel-like extracellular matrix (ECM) proteins (18). Since inhaled particles are mostly accumulated in the ECM (17), there is no reason to ignore the impact of ECM and only focus on the interaction between particles and cells. Thus, we suspected that the lung tissue ECM, apart from the cells, could be also affected by the particles. Indeed, ECM abnormality is closely correlated with cancer development. On one hand, the clinical samples showed the developed tumor was surrounded by dense collagen matrix; on the other hand, one typical example is that tissue fibrosis often precipitates tumor development in the same tissue – such as liver cirrhosis progressing into hepatoma and pulmonary fibrosis leading to lung cancer (14; 52). These clinical and experimental proved reports prompt us to investigate the pathological change of FPM-stimulated lung interstitial ECM and its role in lung tumorigenesis.

Our experimental findings validated this assumption, by demonstrating – and elucidating the mechanism of – FPM-induced alteration in the lung interstitial structure. As soon as we, for the first time, found that FPM in the lung homogenate (LH) dramatically enhanced the crosslinking of soluble type IV collagen (Col IV), we speculated that a specific biomolecule in LH enriched on the FPM surface mediates this action. With proteomics tools, we identified peroxidasin (PXDN); this enzyme, specialized in catalyzing collagen network formation (9), is enriched in the protein corona of FPM, enabling FPM to mediate ECM remodeling. Meanwhile, for PXDN, its adsorption onto FPM triggered its aberrant enzymatic activity through a ‘phase-transition’ process, which further disturbs Col IV crosslinking and the organization of the lung interstitial space. Pre-treatment using PXDN inhibitor could alleviate the tumor-promoting effect of FPM. Therefore, an aberrant ECM remodeling mediated by enriched PXDN in the particle corona is the key pathological change caused by FPM.

Our study further elucidates how such a change in ECM accelerates tumor development. As a consequence of FPM treatment, the denser Col IV and more compacted interstitial space in lung tissue would hinder the migration of CTLs, which are the most ‘informed’ defender to protect against potential cancer and delay malignant progression (15; 32). Once CTLs’ migration track was blocked, the CTLs infiltration would be impaired, thus decreasing the chance of cell-cell cytotoxic interaction to tumor cells in lung tissue (25). The CTLs’ insufficient early response to tumor cells indicated limited immune surveillance and considerably dropped tumor recognition. All the evidences recapitulate how FPM accelerated the development of lung cancer through interfering the lung immune microenvironment, shedding light on the sequential consequences between particle-induced ECM remodeling, impaired immunosurveillance, and tumorigenesis.

Taken together, our study reveals a completely new mechanism by which inhaled fine particles promotes lung tumor development. This mechanism is notable in three aspects. First, we herein provide direct evidence that protein corona on those foreign objects can elicit such a significant and adverse effect. Although corona formation on nanoparticles has been extensively studied in recent years, its involvement in a pathogenic process related to a global health issue is rare. Our findings here highlight the importance of the corona-endowed, ‘new’ bioactivity of nanomaterials *in vivo* – and even in a particular tissue. Second, PXDN, or other proteins mainly engaged in ECM modulation, is less expected as a major player in lung carcinogenesis, especially at the initial stage, until this study reveals it as a specific and unexpected molecular target for FPM. These investigations enable the specific design of PXDN-targeted preventive or therapeutic approaches. The relationship between the physicochemical properties of FPM and the affected PXDN activity should be further explored in greater detail. Third, in a specific organ (the lung), our data demonstrate that physical blockage of immunocytes movement directly increases tumorigenesis, which suggests an important previously unconsidered role of the *in vivo* delivering or deploying the power of the immune system in various immune-oncology processes.

## Materials and Methods

### FPM Collection and Preparation

Fine Particulate Matter (FPM) samples within sizes of 1 μm was collected using TH-16A multiple-channel atmospheric particulate automatic sampler (Wuhan Tianhong Instrument Ltd, Wuhan, China) and filtered them through Whatman PTFE membranes (GE Healthcare Life Sciences, Pittsburgh, USA). For particles in air pollutants (PM1), samples were conducted continuously for 7 days at different representative of Nanjing City (Qixia, Jiangning, Pukou, Gulou, and Gaochun) in Jiangsu Province, Suzhou City in Anhui Province and Tieling City in Liaoning Province (named as: QX, JN, PK, GL, GC, SZ and TL). For TSP samples, the residual smoke of burned tobaccos with filters were collected in customized confined space. Then PTFE filter membranes containing FPM were cut into 0.1 cm × 0.1 cm pieces, immersed in distilled water for 2 days and oscillated ultrasonically for 1 h for 3 times. Detached FPM was separated with filter membranes after centrifugation at 2,000 rpm for 5 min for 3 times. Supernatant enriched FPM were vacuum freeze-dried and then stored at – 20 °C. For preparation of FPM suspension, FPM samples were suspended in sterile 1× PBS (phosphate buffer saline) to achieve 10 mg/mL particles for further analysis.

### Establishment and Treatment of Lung Tumor Model in FPM-exposed Mice

#### FPM-exposed mice

Mice exposed to FPM was generated according to a previously published method (57). six-to-eight weeks old C57BL/6 mice or *Foxn1^nu^* naked mice of the same ground were purchased from Beijing Vital River Laboratory Animal Technology Co. Ltd. (Beijing, China). OT-1 T cell receptor transgenic mice [C57BL/6-Tg (TcraTcrb) 1100Mjb/J] were a gift from K. Zeng (Nanjing University, China). These mice were randomly divided into 3 groups (PBS ones, FPM-exposed ones: PM1 and TSP; each group contained at least 3 mice). Mice were anesthetized by intraperitoneal injection of pentobarbital sodium at 45 mg/kg body weight. After the trachea exposed by opening the neck skin and blunt dissection, mice received suspension of 0.2 mg FPM in a total volume of 50 μl of sterile physiological saline by inserting a 7-gauge needle (BD Biosciences, San Jose, CA, USA) into the trachea trans-orally. To be estimated, before intratracheal instillation, FPM suspension was always sonicated and vortexed. After the site of surgery sutured and cleaned with penicillin, the mice were allowed to recover until they were sacrificed. As control, PBS was applied in a similar manner. After exposed to FPM for 7 days, mice were 1) sacrificed for analyzing the changes of lung tissue structure, or 2) subsequently stimulated with 5 μg/kg T cell chemokine – C-X-C motif chemokine ligand 10 (CXCL10, PeproTech, Rocky Hill, USA) (23), or named as interferon-inducible protein-10 (IP-10) for 2 h through intratracheal injection, to analyze the CTLs’ infiltration into lung tissue, 3) to analyze the location of PXDN and FPM in the lung, mice were exposed to rhodamine-labelled FPM (R-FPM) with intraperitoneal injection. Lung tissue was extracted for immunofluorescence 4 hours later.

#### Establishment of lung tumor model in FPM-exposed mice

For the syngeneic LLC (Lewis lung carcinoma) model, after exposed to FPM for 7 days, mice were further intravenously injected with 5×10^7^ LLC cells for indicated days (0 d, 1 d, 3 d, 5 d, 10 d and 20 d) to create the lung carcinoma model post particle administration. For the LLC-stimulated carcinoma model, 20 days after the injection of LLC, the mice of different groups were sacrificed, and the lung tissue were extracted for analysis. The number of tumors suffered by the mice was examined and evaluated randomly under blindfold conditions. For the transgenic (K-ras^G12D^p53^−/−^) mouse models, mice harboring a Cre-inducible endogenous oncogenic *Kras^LSL-G12D^p53^fl/fl^* allele (GemPharmatech Co.,Ltd, Nanjing, China), were treated as above. 7 days after exposed to FPM, mice were allowed to inhale 1×10^7^ plaque forming units (PFU) Cre adenovirus (AdCre, OBiO Technology (Shanghai) Corp., Ltd., Shanghai, China) to activate K-ras^G12D^ expression and knock out p53 in lung tissue (*Kras^G12D^p53^−/−^* transgenic model). The mice were sacrificed 50 days after tumor initiation.

#### Administration of PXDN inhibitor

A series of plasmids capable of ectopically expressing PXDN specific short hairpin RNA (shPXDN) or control shRNA (shNC) were designed and constructed by Genepharma Biotechnology (Shanghai, China). For transfection, *in vivo*-jetPEI (Polyplus Transfection, Illkirch, France) was used as a delivery agent (20). The transfection reagent complex (0.16 μL of *in vivo*-jetPEI per μg plasmid DNA) was prepared and mixed according to the manufacturer’s instructions in glucose buffer. Then the 20 μl buffer containing 4 μg shPXDN was delivered into murine lung tissue through trachea injection. Besides, to analyze the effect of small molecular PXDN inhibitorS, the FPM-exposed mice were administrated with 25 mg/kg methimazole (MMZ, MedChemExpress LLC, Shanghai, China) and 50 mg/kg phloroglucinol (PHG, Aladdin, Shanghai, China) 3 days pre- and post-FPM stimulation respectively.

During the experimental procedure, all animal studies were performed under protocols approved by institutional guidelines (Nanjing University Institutional Animal Care and Use Committee). They were also required to be conformed to the Guidelines for the Care and Use of Laboratory Animals published by the National Institutes of Health. The mice were housed 5 per cage and fed in a specific pathogen-free (SPF) animal facility with controlled light (12-h light/dark cycles), temperature and humidity, with food and water available.

### Analysis of T Cell Migration on Lung Tissue

#### Lung tissue slice preparation

To analyze T cell migration on lung tissue, the lung samples of different group, including the FPM-exposed mice and the FPM-exposed mice pre-treated with PXDN shRNA were respectively prepared as the 50 μm frozen section (46). In some experiments, tissue sections were pre-treated with 50 μg/mL collagenase D (Worthington Biochemical Corp., Colorado, USA) in RPMI 1640 for 5 min, then rinsed in complete RPMI 1640 medium.

#### Cell preparation

Jurkat T cells were stained with Calcein-AM (DOJINDO LABORATORIES) for 30 minutes at 37°C and then washed with HBSS (Sangon Biotech) for three times. 1.5 × 10^5^ T cells total in 10 – 20 μl were added at one side of the cut surface of each slice. To ensure cells settle down on the slice, slices with T cells were incubated for 1 hour at 37°C, 5% CO_2_; gently washed to remove the residual cells that had not entered the tissue; and kept at 37°C, 5% CO_2_ before imaging.

#### Time-lapse imaging and cell trajectories analysis

For imaging T cells’ migration on the lung tissue slice, 5 μg/mL IP-10 were added on the other side of the slice and images were then acquired in time-lapse model with a SP5 confocal system (Leica) every 3 min for 1 h. Imaging was exported and compressed into videos as avi. format. To quantify T cell trajectories with the surrounding ECM in lung tissue, the cell migration video including image-sequences cell migration was analyzed with TimTaxis Software (https://www.wimasis.com/en/WimTaxis) by identification of the centroids of individual cells at consecutive timepoints. The relationship of statistical data composed of displacement, distance, velocity and acceleration *vs.* time were respectively further analyzed.

### Mass Spectrometry and Identification of Sulfilimine Bond Crosslinked Peptides

To analyze effect of FPM on crosslink of collagen IV, the potential reaction site containing sulfilimine bond (NC1 domain formed along with C-terminal aggregation) was detected with liquid chromatography-mass spectrometry (LC-MS) (9; 38). Briefly, 5 mg/mL soluble and commercially available collagen IV (Sigma-Aldrich, St. Louis, MO, USA), which was extracted from murine sarcoma basement membrane, was incubated with LH or LH-FPM mixture, 100 μM H_2_O_2_ and 200 μM NaBr for 4 h at 37 °C. After centrifugation at 12,000 rpm for 20 min, the crosslink pellet was collected. Then the pellet was digested with collagenase D (50 μg/mL; Worthington) for 30 min at 37 °C to yield the peptide containing the crosslink site. After centrifugation at 12,000 rpm for 20 min, the supernatant containing the crosslinked peptides was collected. After separation by SDS-PAGE, NC1 domain was digested with trypsin (MS Grade, Thermo Fisher Scientific) overnight at 37 °C and then analyzed by LC-MS analysis on a Shimadzu UFLC 20ADXR HPLC system in-line with an AB Sciex 5600 Triple TOF mass spectrometer (AB SCIEX, Framingham, Massachusetts State, USA). To analyze low abundance peptides containing crosslinked domain, targeted methods were performed with PeakView software (AB SCIEX) based on raw continuum LC-MS data. Briefly, full scan spectra of total ion chromatography (TIC) diagram were acquired, and LC-MS peptide reconstruct with peak finding were provided. According to calculated theoretic mono-isotopic mass of the sulfilimine (the mass of two hydrogen atoms was subtracted from the sum of the masses for Met93-containing peptide and Lys211-containing peptide), corresponding mass spectrum (about 5030.425, 5046.425 or 5062.425) with different oxidation of methionine (M_ox_) were specifically searched. To delineate the difference of crosslink site, the extract ion chromatography (XIC) diagram based on corresponding was displayed and compared.

### Analysis of Peroxidasin Enzyme Activity and Reaction

#### Measurement of peroxidasin activity

To analyze effect of FPM on PXDN, the enzyme activity was analyzed by Amplex Red hydrogen peroxidase assay kit (43) (Thermo Scientific). Briefly, after incubated with FPM for 30 min at 37 °C, the enzyme activity was detected in reaction containing 50 mM Amplex Red reagent, 1 mM H_2_O_2,_ and PXDN (PeproTech) mixed in FPM. After incubation for 30 min at room temperature, fluorescence was measured at excitation wavelength 590 nm with a fluorescence microplate reader (Thermo Fisher Scientific, MA, USA). To avoid interference of particles per se, the equal FPM was set as negative control.

#### Analysis of peroxidasin enzyme kinetics

To detect the change of PXDN catalytic efficiency after its incubation with FPM, PXDN enzyme kinetic behaviors were investigated according to the Michaelis–Menten model. Briefly, after incubated with FPM for 30 min at 37 °C, 10 mU PXDN was mixed with a serial concentration of H_2_O_2_ (0, 10, 50, 100, 150, 200 and 250 μM). The enzymatic reaction was analyzed by adding 50 mM Amplex Red detection reagent (35). The fluorescent reaction was measured every 40 sec for 8 min and subsequent every 5 min for 6 times using a microplate reader (Thermo Scientific) with enzyme kinetics model. Line weaver– Burk representative plot was generated from the relationship of reaction rate and substrate concentration. Based on the plot, the Michaelis constant (Km) reflecting the binding efficiency of the enzyme with the substrate and turnover number (Kcat).

#### Detection of hypobromous acid

Based on that NADH could be stably brominated into NADH bromohydrin after its reaction with hypobromous acid, hypobromous acid generated by PXDN and its bromination activity were tested by TripleTOF^TM^ 4600 LC-MS/MS according to the reported literature with a little modification (6). Briefly, 100 nM PXDN or the mixture of PXDN and FPM was incubated at 37 °C in PBS containing 200 μM NADH, and 200 μM NaBr for 30 min. Reactions were started upon addition of 100 μM H_2_O_2_. Then the supernatant was separated and analyzed to detect NADH and its bromohydrin products. NADH was measured using the transition m/z 664.2 to 408.1, and the bromohydrins by m/z 760.2 and 762.2 both going to m/z 680.2. Intensity of peaks were calculated using PeakView software (AB SCIEX). Then the ration of NADH bromohydrin relative to NADH based their peak intensity was calculated.

#### Measurement of hypochlorous acid

Reactions were initiated with the addition of 100 μM H_2_O_2_ and 200 μM NaCl after 1 μg PXDN incubated with FPM for 30 min. 5 μM HClO-detecting fluorescent probes (kindly provided by ICMS) were added to react for 30 min and fluorescence intensity was determined at excitation wavelength 488 nm by a fluorescence microplate reader (62). Then HOCl production was calculated according to the standard curve of a serial concentration of HOCl *vs.* absorbance. To be estimated, HOCl standards should be freshly prepared by adjusting the pH of NaClO to 7.4 to create HOCl solutions before each time.

#### Measurement of NC1 crosslink by peroxidasin

To delineate sulfilimine crosslink of NC1 fragment in the collagen IV, the solubilized NC1 monomer purified from mouse renal basement membrane (Chondrex, Inc., WA, USA) was incubated with PXDN or the mixture of PXDN and FPM. To initiate the reaction, 100 μM H_2_O_2_ and 200 μM NaBr was respectively added. After incubation for 30 min at 37 °C, to visualize the change of sulfilimine crosslinked dimeric (NC1_di_) and non-crosslinked monomeric subunits (NC1_mo_), the solution underwent sodium dodecyl sulfate polyacrylamide gel electrophoresis (SDS-PAGE) under non-reducing conditions followed by Coomassie Blue staining.

### Identification and Characterization of PXDN’s Liquid-Liquid Phase Separation

#### Microscopy analysis of liquid-liquid phase separation (LLPS)

To analyze the droplet formation of PXDN under the stimulation of FPM, 10 mM fluorescein isothiocyanate (FITC) labelled proteins were incubated with 50 μg/mL rhodamine B labelled FPM for 30 min in LH. Then samples of different group were dropped onto a glass slide and sealed with a coverslip. Phase separation of PXDN and its liquid-like droplets was observed under phase contrast and confocal microscopy with a 100x Oil objective (Nikon). The distribution profile of fluorescence intensity of liquid-like PXDN and FPM were respectively analyzed by the Nikon NIS-Elements software. Besides, to predict the domain that might trigger PXDN’s accumulation, the intrinsically disordered regions (IDR), the domain frequently closed to proteins’ phase-separation, of PXDN is predicted based on the IUPred algorithm (https://iupred2a.elte.hu/).

#### Fluorescence recovery after photobleaching (FRAP) assays

After 10 mM fluorescein isothiocyanate (FITC) labelled proteins were incubated with 50 μg/mL rhodamine B labelled FPM for 30 min in LH, fluorescence recovery after photobleaching experiments were performed on rhodamine-labelled PXDN droplets formed in PM1. The photorecovery behavior was tracked using the 564 nm laser line of a 40 × 1.0NA objective on Zeiss LSM 980 with 2.4-fold magnification. Photobleaching was done with 100% laser power to 30% intensity using the bleaching program of the ZEN software and time-lapse images were recorded every 10 s. After bleaching, the fluorescence intensities were measured and collected by mean ROI (photo-bleached region and control region without bleach). The raw data with three bleach treatment are processed and analyzed by GraphPad Prism.

### Molecular Docking on The Effect of Phase Separation on Enzymatic Reaction

Template crystal structures of PXDN and NC1 domain in Col IV were identified and downloaded from Swiss Model Protein Data Bank as the PDB files (PXDN: 5MFA.1A; NC1: 5NAY). Besides, the putative structure under phase transition, which simulated the PXDN assembly, was created through homology modeling based on one experimentally determined structures of peroxidasin-related family member (PDB ID: 4C1M) in the RCSB Protein Data Bank (8). Subsequently, the most similar template conformation with 49.8% consistency was chosen from among the candidates and named as the PXDN Dimer. Then ZDock protocol was used for molecular docking analysis of the interaction between PXDN and NC1. Docking models of intuitive contacting interface was outputted after scoring and selection. The interactive area was especially labelled.

### Statistical analysis

The results are expressed as mean ± standard deviation (SD). Data were statistically analyzed using Prism software (GraphPad) and assessed for normality or homogeneity of variance. Differences between multiple groups were compared using one-way or two-way analysis of variance (ANOVA) with Dunnett’s tests or, if appropriate, repeated measures ANOVA test with *post-hoc* Bonferroni correction. Differences between two groups were evaluated using the two-tailed unpaired Student’s t-test. A value of *p* < 0.05 was considered significant; n.s. = not significant.

## Supporting information

Supplementary Video S1_Lung Tissue_PBS

Supplementary Video S2_ Lung Tissue_PM1

Supplementary Video S3_ Lung Tissue_TSP

Supplementary Video S5_PM 1-Collagenase

Supplementary Video S6_TSP

Supplementary Video S7_TSP-Collagenase

## Data Availability

The data generated in this study are available within the article and its supplementary data files. Additional data related to this paper are available upon request from the corresponding author.

## Author Contributions

Z.W. and L.D. conceived, performed, and analyzed all experiments and wrote the manuscript. J.Z. designed the study. C.W. designed the study and wrote the manuscript. Z.Z., C.C., X.T., Z.X. and P.X. performed the experiments. Y.Y. performed data analysis and molecular docking.

## Acknowledgements

We thank Professor Ke Zeng in Nanjing University for his kindly providing the OT-1 TCR transgenic mice and OVA-LLC cells. This study was funded by the National Natural Science Foundation of China (31971309, 31671031, 32001069, 81973273, 81673380), the Natural Science Foundation of Jiangsu Province (BK20200318), and the Fundamental Research Funds for the Central Universities (020814380088, 020814380115). C.W. acknowledges the financial support from the Science and Technology Development Fund, Macao SAR (FDCT 0097/2019/A2, 0018/2019/AFJ) and the University of Macau Research Committee (MYRG2019-00080-ICMS). This study also supported by the funds for the International Cooperation and Exchange of the Natural Science Foundation of China and the Science and Technology Development Fund (31961160701).

