## Supplementary Information for "Air Pollution Particles Hijack Peroxidasin to Disrupt Immunosurveillance and Promote Lung Cancer"

1. **Characterization of FPM.**

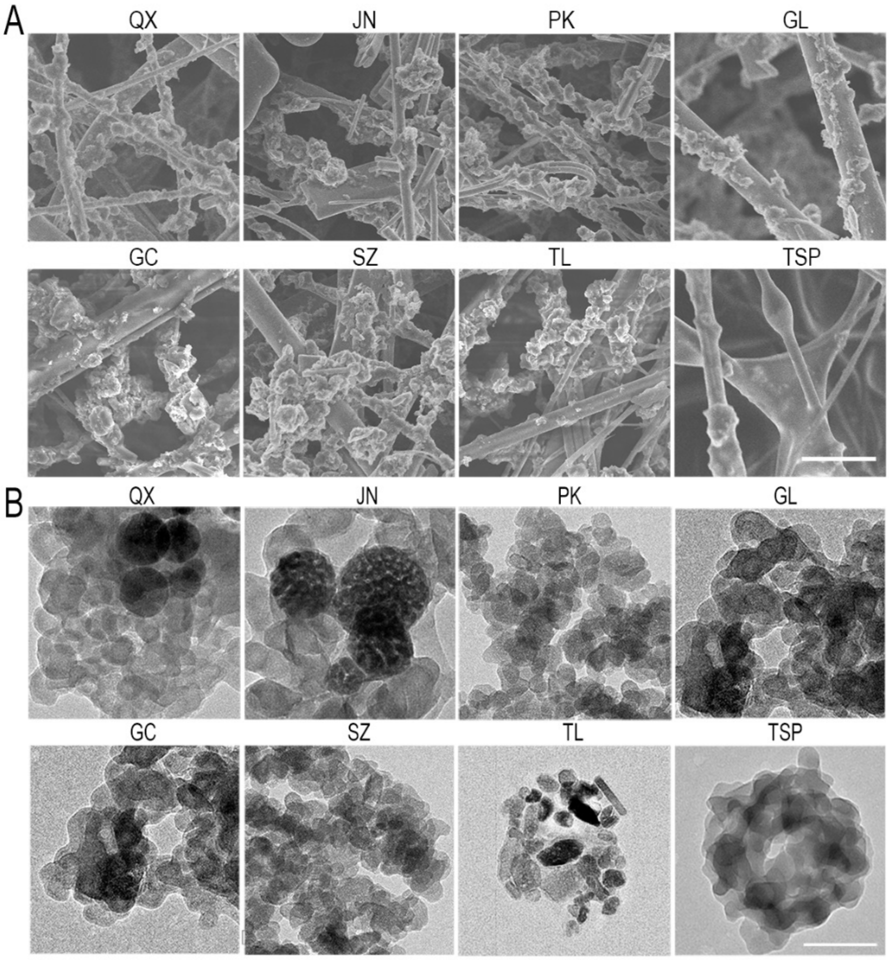

**Supplementary Figure S1.** A. Representative scanning electronic microscope (SEM) for particles detained on the fiber filter membrane, which were collected from airborne pollution in 7 different locations, including Nanjing City (Qixia, Jiangning, Pukou, Gulou, and Gaochun) in Jiangsu Province, Suzhou City in Anhui Province and Tieling City in Liaoning Province (named as: QX, JN, PK, GL, GC, SZ and TL) and tobacco smoke particle (TSP). Scale bar = 5 μm; B. Representative transmission electron microscopy (TEM) images for detached particles from filter membrane in panel A. Scale bar = 100 nm. Images are representative for three independent experiments.

**Supplementary Table S1**. Physicochemical characteristic of fine particulate matter collected from airborne pollution in 7 different locations, including Nanjing City (Qixia, Jiangning, Pukou, Gulou, and Gaochun) in Jiangsu Province, Suzhou City in Anhui Province and Tieling City in Liaoning Province (named as: QX, JN, PK, GL, GC, SZ and TL) and tobacco smoke particle (TSP), including the size distribution, zeta potential and the top 15 element compositions.

| | QX | JN | PK | GL | GC | SZ | TL | TSP |
|---|---|---|---|---|---|---|---|---|
| Size (nm) | 207.6±93.4 | 185.0±68.3 | 208.1±112.3 | 206.7±77.8 | 214.7±90.7 | 180.0±88.1 | 203.8±82.4 | 216.5±83.6 |
| Potential (mV) | - (29.53±2.97) | - (29.77±0.25) | - (35.1±2.17) | - (36.7±1.15) | - (39.23±0.45) | - (35.1±2.17) | - (37.73±0.71) | - (16.3±1.59) |
| Top 15 Element Compositions (%) | N (16.8100) | N (12.2000) | N (18.5900) | Ca (13.6875) | C (8.8390) | C (8.8420) | C (23.1900) | C (48.7700) |
| | Ca (14.625) | C (8.6310) | C (11.9600) | N (11.8800) | N (6.4650) | N (3.4100) | S (5.8280) | N (6.9490) |
| | Al (8.6875) | H (4.3630) | S (6.5530) | C (9.3640) | Ca (4.6422) | Ca (3.0000) | N (5.2780) | H (6.9240) |
| | C (7.7410) | S (4.2000) | H (6.4900) | Al (5.1875) | H (4.3440) | S (2.4500) | H (4.5260) | Ca (2.1837) |
| | S (6.1480) | Ca (3.3000) | Ca (6.4438) | S (4.0600) | S (4.3410) | H (2.2150) | Ca (1.5625) | S (1.1760) |
| | H (5.5500) | Al (0.9375) | B (1.1880) | H (3.8680) | Al (3.7203) | K (1.1563) | K (1.5250) | Al (0.2218) |
| | Mg (3.0625) | Fe (0.4181) | Zn (1.0438) | B (2.0938) | B (1.6016) | Ba (0.1813) | B (0.3938) | B (0.1663) |
| | B (3.0000) | K (0.3438) | K (0.7438) | Mg (1.6250) | Mg (1.1470) | Cu (0.1266) | Zn (0.3375) | K (0.1365) |
| | Na (1.1188) | B (0.2125) | Na (0.7188) | K (0.8625) | K (0.7375) | Zn (0.1000) | Al (0.2563) | Mg (0.0921) |
| | K (1.0000) | Zn (0.1125) | Fe (0.3843) | Fe (0.3938) | Fe (0.4788) | Al (0.0750) | Ba (0.2438) | Zn (0.0145) |
| | Fe (0.7188) | Pb (0.0363) | Cu (0.2211) | Zn (0.2438) | Zn (0.4594) | Fe (0.0575) | Cu (0.1179) | Cu (0.0067) |
| | Zn (0.4375) | Mg (0.0356) | Ba (0.1063) | Na (0.1375) | Na (0.3172) | Pb (0.0231) | Fe (0.1125) | Cd (0.0039) |
| | Ba (0.3938) | Sr (0.0331) | Al (0.0688) | As (0.0863) | Cu (0.1004) | Ti (0.0201) | Na (0.1000) | Sr (0.0034) |
| | As (0.1425) | Mn (0.0258) | Pb (0.0519) | Pb (0.0744) | As (0.0572) | Mn (0.0102) | Ti (0.0388) | As (0.0031) |
| | Cu (0.0934) | Cu (0.0133) | Mn (0.0252) | Cu (0.0738) | Pb (0.0520) | Sr (0.0100) | Pb (0.0231) | Fe (0.0026) |

### 2. Effect of FPM on Lung Tumorigenesis.

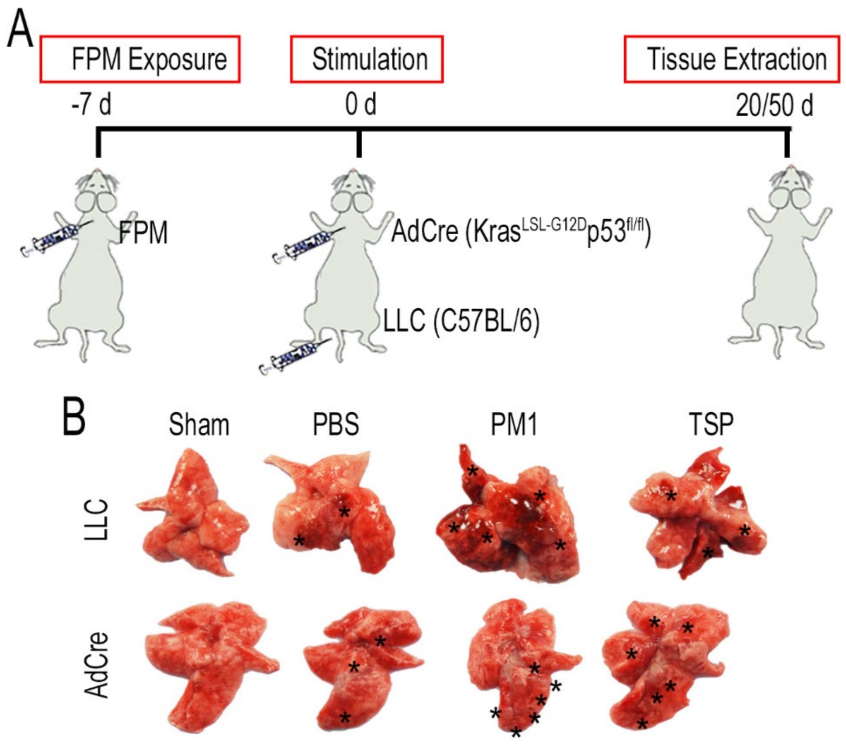

**Supplementary Figure S2. A.** Schematic diagram of Lewis lung carcinoma (LLC)-stimulated or Kras$^{G12D}$/p53$^{-/-}$-transgenic lung cancer model with FPM exposure; **B.** Gross lung tissue images in the FPM-exposed mice of LLC model or Kras$^{G12D}$/p53$^{-/-}$- model 20 days or 50 days after mice were stimulated with LLC or Cre-inducible adevirus (AdCre). The tumor sites were labelled with black asterisk.

### 3. Effect of FPM on Tumor Cells or Cytotoxic T Lymphocytes.

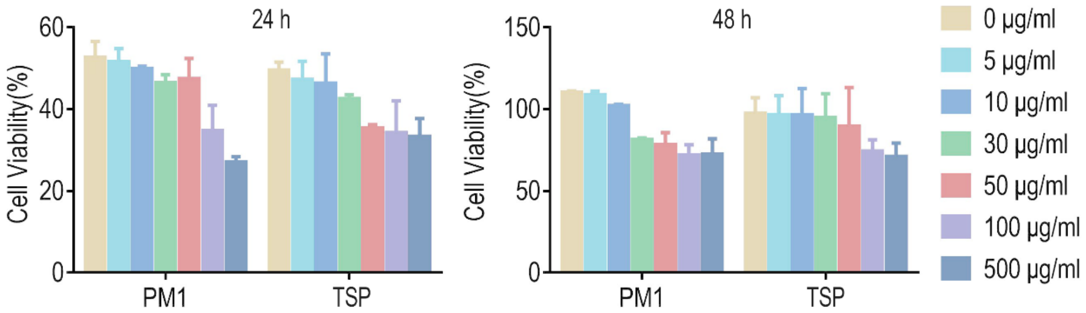

**Supplementary Figure S3.** Cell cytotoxic analysis of LLC cells stimulated with a serial concentration of FPM (0, 5, 10, 30, 50, 100 and 500 μg/mL) for 24 h and 48 h. n=3. Results are shown as mean ± SD.

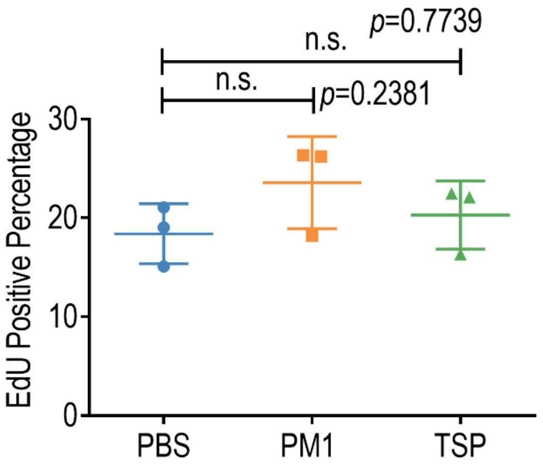

**Supplementary Figure S4.** The quantified analysis of EdU-positive cells in tumor site of lung tissue of LLC model 20 days after tumor initiation. n=3. Results are shown as mean ± SD. n.s. indicates no statistical significance.

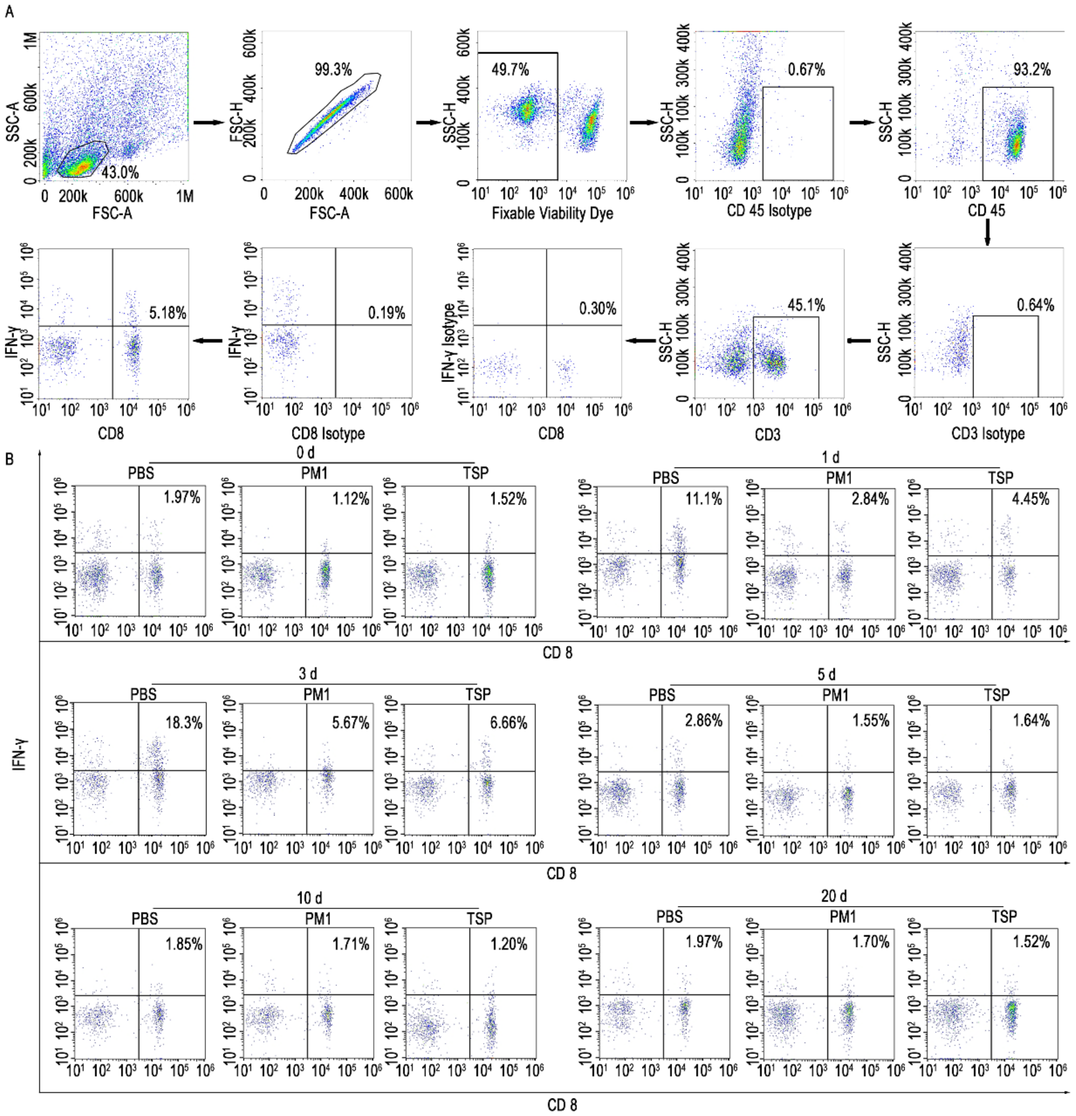

**Supplementary Figure S5. A.** Gating strategy for the quantification of cytotoxic T lymphocytes CTLs cells (IFN-$\gamma^{+}$CD8$^{+/}$CD45$^{+}$CD3$^{+}$) in the lung tissues. **B.** Representative flow cytometry analysis of CTLs in FPM-exposed lung tissue of mice under the physical conditions (0 d) or at indicated days (0d, 1 d, 3d, 5 d, 10 d and 20 d) after intravenous injection with LLC.

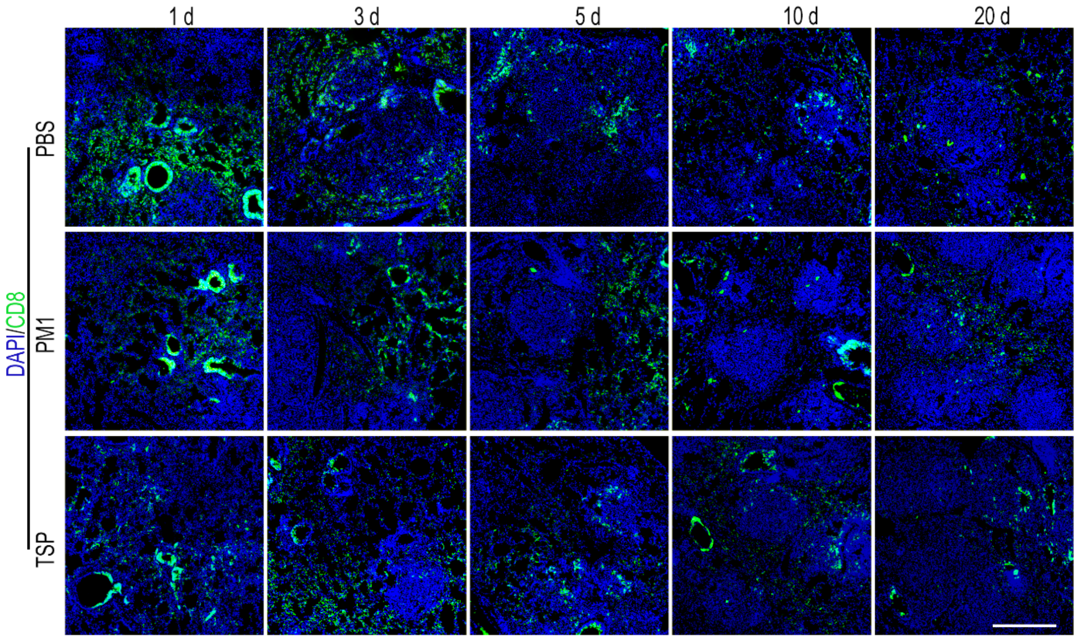

**Supplementary Figure S6.** A. Representative immunofluorescence images of CTLs' infiltration into the FPM-exposed lung tissue at indicated days (1 d, 3 d, 5 d, 10 d and 20 d) after intravenous injection with LLC. The CTLs were labelled with CD8 and shown in green. Images are representative for three independent experiments. Scale bar = 200 µm.

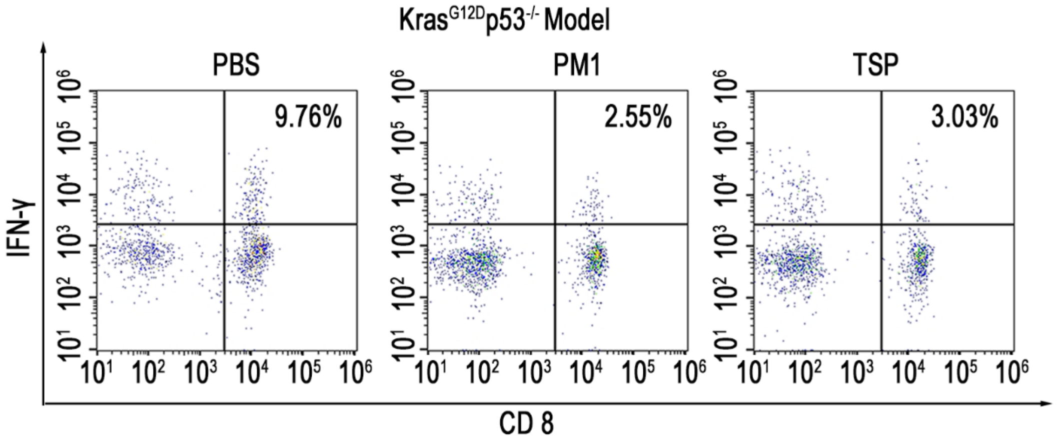

**Supplementary Figure S7.** Representative flow cytometry analysis of CTLs (IFN-$\gamma^{+}$CD8$^{+}$/CD45$^{+}$CD3$^{+}$) in FPM-exposed lung tissue of K-ras$^{G12D}$p53$^{-/-}$ mice 4 weeks after tumor initiation with the intratracheal injection of AdCre.

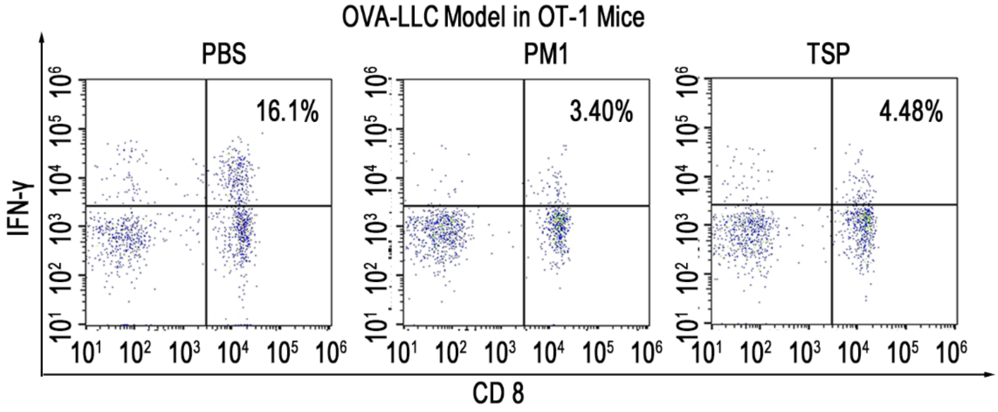

**Supplementary Figure S8.** Representative flow cytometry analysis of CTLs (IFN-$\gamma^{+}$CD8$^{+}$/CD45$^{+}$CD3$^{+}$) in FPM-exposed lung tissue of OT-1 TCR transgenic mice 1 day after OVA-LLC stimulation.

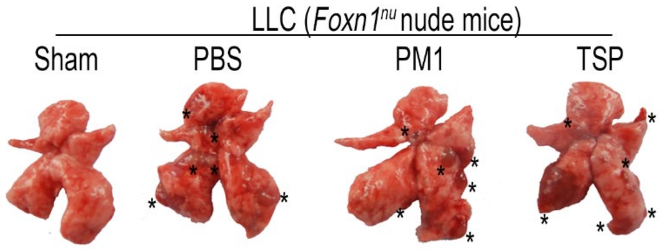

**Supplementary Figure S9.** Gross lung tissue images in FPM-exposed *Foxn1$^{nu}$* nude mice 20 days after they were intravenously injected with LLC.

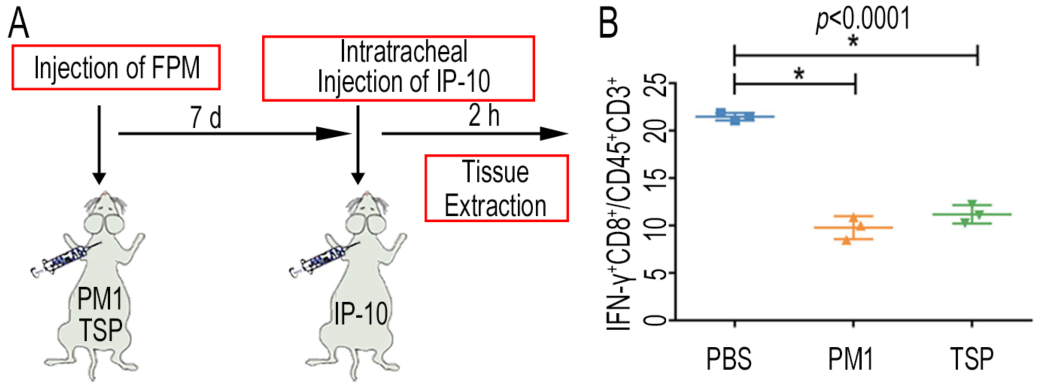

**Supplementary Figure S10.** A. Scheme of analyzing the effect of T cell chemokines IP-10 on the CTLs' infiltration into the FPM-exposed mice; B. Statistical flow cytometry analysis of CTLs cells (IFN-$\gamma^{+}$CD8$^{+}$/CD45$^{+}$CD3$^{+}$) in lung tissue 2 hours after the mice were stimulated with 5 μg/kg IP-10 through intratracheal injection. n=3. Results are shown as mean ± SD. *$p<0.05$ after ANOVA with Dunnett's tests.

### 4. Effect of FPM on CTLs' Migration Potential.

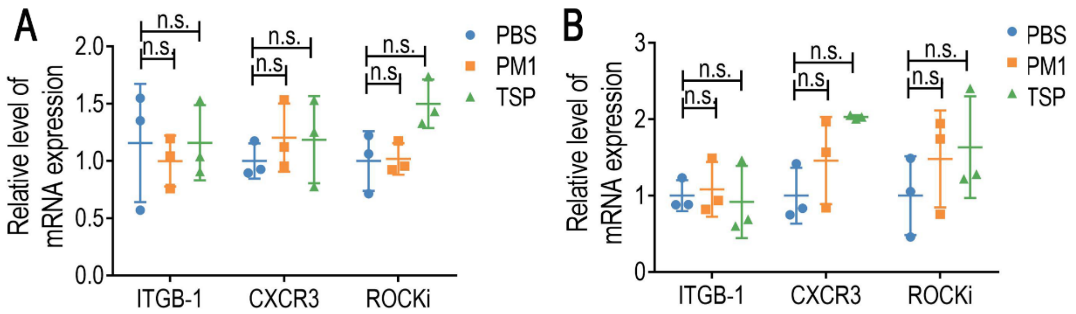

| ITGB-1 | |
|---|---|
| PBS vs. PM1 | 0.1578 |
| PBS vs. TSP | -0.001037 |
| CXCR3 | |
| PBS vs. PM1 | -0.204 |
| PBS vs. TSP | -0.1866 |
| ROCKi | |
| PBS vs. PM1 | -0.01883 |
| PBS vs. TSP | -0.4998 |

| ITGB-1 | |
|---|---|
| PBS vs. PM1 | -0.08206 |
| PBS vs. CSP | 0.08206 |
| CXCR3 | |
| PBS vs. PM1 | -0.4599 |
| PBS vs. CSP | -1.031 |
| ROCKi | |
| PBS vs. PM1 | -0.4811 |
| PBS vs. CSP | -0.6348 |

**Supplementary Figure S11.** Transcriptional level of typical markers related to CTLs' migration, integrin-1 (ITGB-1), C-X-C motif chemokine receptor 3 (CXCR 3) and Rho-associated kinase (ROCKi), in Jurkat T cells after they were stimulated with FPM for 48 h (A) or in the CTLs separated from lung tissue exposed to FPM for 7 days (B). n=3. Results are shown as mean ± SD. n.s. indicates no statistical significance. *p* values were listed underneath.

### 5. Effect of FPM on Lung Structure.

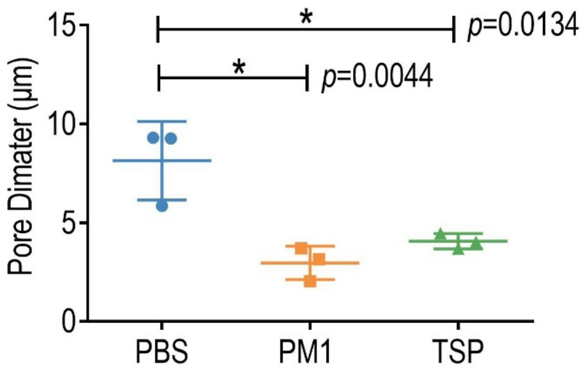

**Supplementary Figure S12.** The quantified analysis of pore diameter of interstitial matrix in the lung tissue, based on the scanning electron microscope (SEM) images and analyzed by Image J. n=3. Results are shown as mean ± SD. **p*<0.05 after ANOVA with Dunnett's tests.

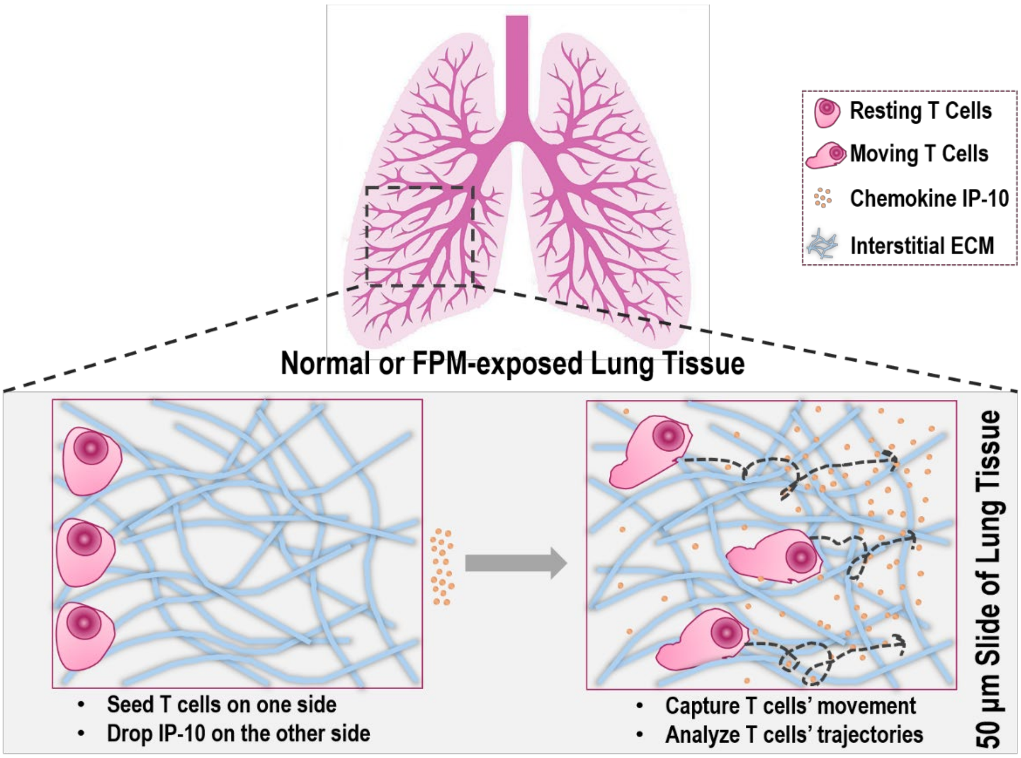

**Supplementary Figure S13.** Schematic diagram of analyzing of CTLs' migration in lung tissue slice of FPM-exposed mice or PBS group.

**Supplementary Video S1-3.** Dynamic migration videos of T cells in the slice of lung tissue exposed to FPM. Jurkat T cells were pre-stained with Calcein-AM and labelled as green in videos.

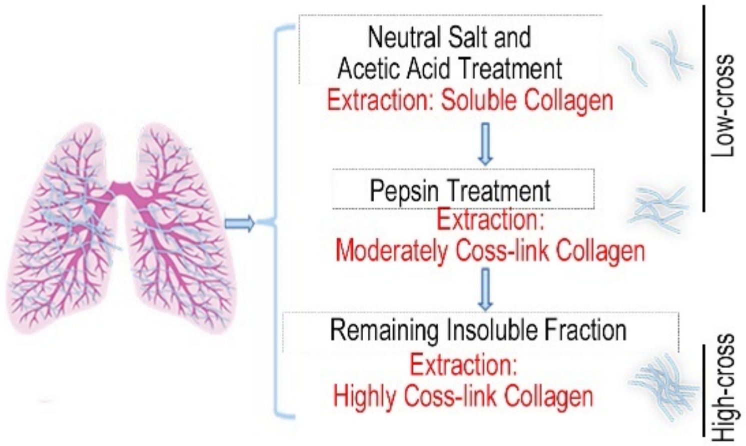

**Supplementary Figure S14**. Schematic diagram of separating collagen fraction with different crosslink level. The part separated by neutral salt, acetic acid and pepsin was defined as 'low-cross' and the residual ones was regarded as 'high-cross', according to the reported literature (4).

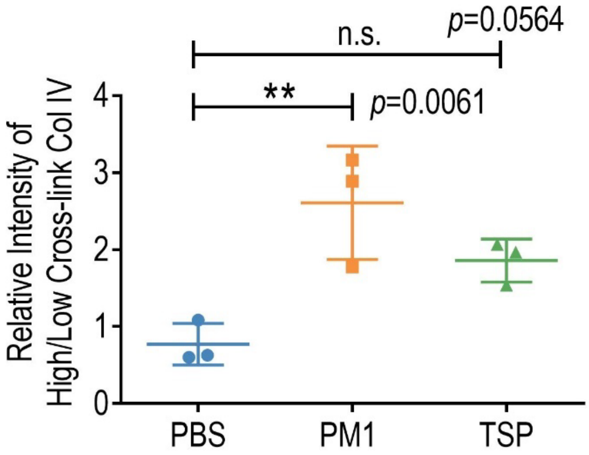

**Supplementary Figure S15.** The relative intensity of high-crosslink Col IV to low-crosslink ones according to the WB results of lung tissue exposed to FPM for 7 days, based on the Image J analysis. n=3. Results are shown as mean ± SD. *$p$<0.05 after ANOVA with Dunnett's tests. n.s. indicates no statistical significance.

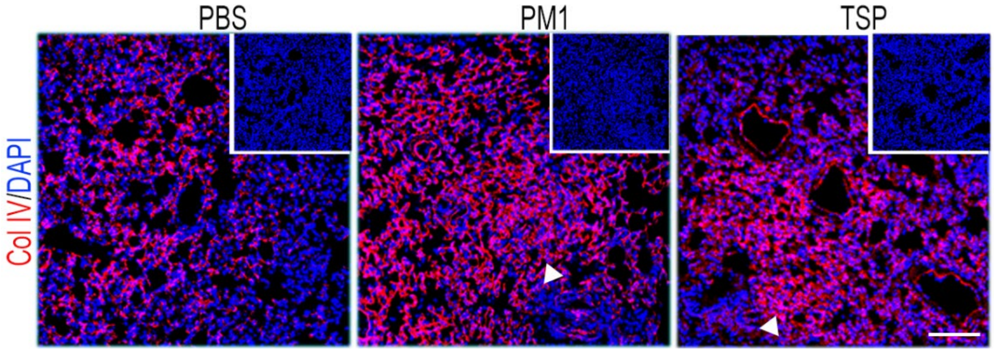

**Supplementary Figure S16.** Representative Col IV immunofluorescence images of lung tissue in the mice exposed to FPM for 7 days, with the blue DAPI staining images shown in the inserted box. Scale bar = 100 μm.

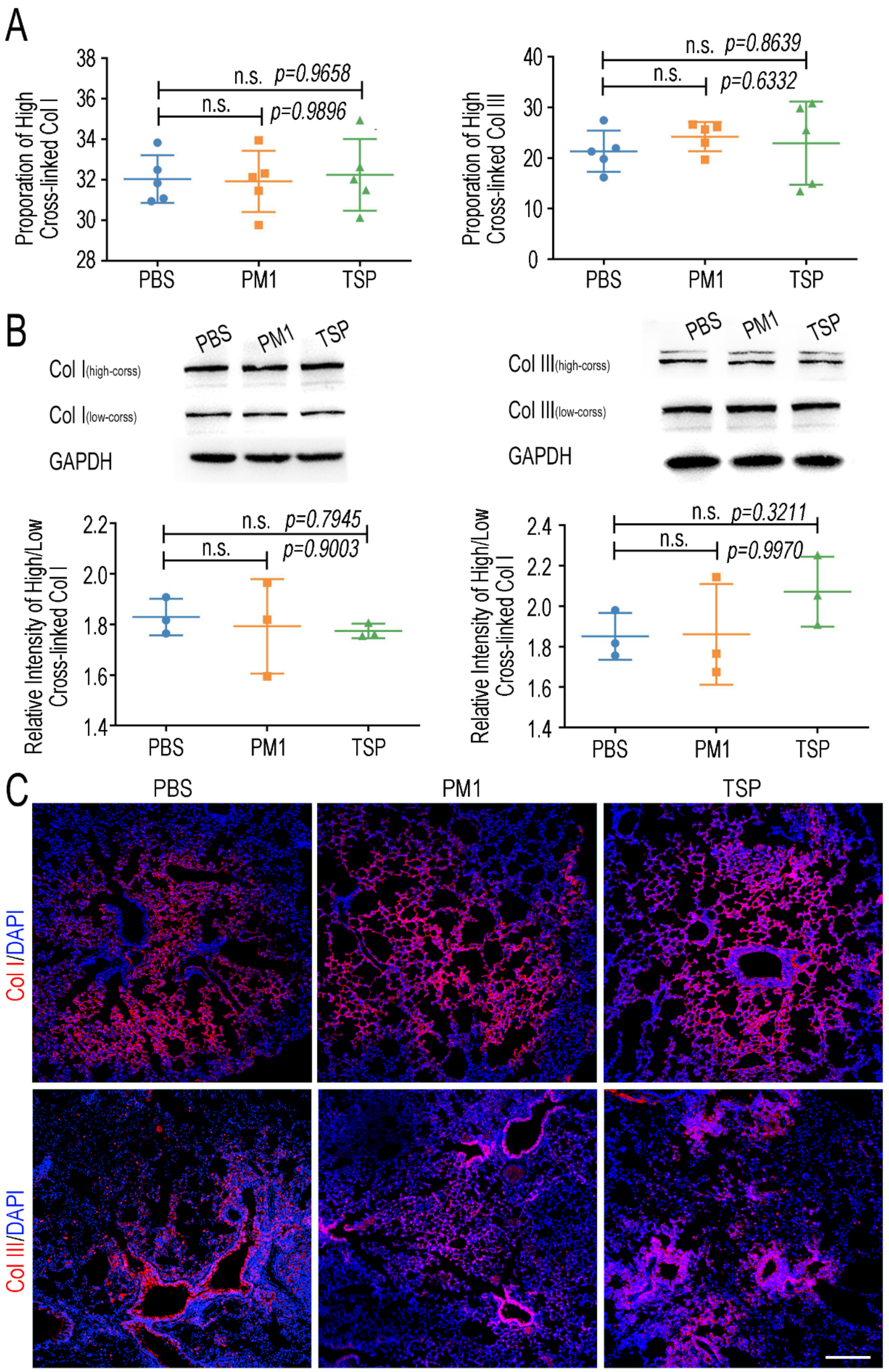

**Supplementary Figure S17.** A. Proportion of high-crosslink Col I (left) and Col III (right) in lung tissue of mice exposed to FPM or PBS for 7 days, which was calculated by the 'high-cross' fragment divided by the sum of different fraction ('low-cross' ones and 'high-cross' ones). The content of each part was detected by enzyme linked immunosorbent assay (ELISA). n=5.; B. Western blotting analysis of 'low-cross' collagen and 'high-cross' Col I (left) and Col III (right) in lung tissue of mice exposed to FPM or PBS for 7 days. Their relative intensity analyzed by Image J was shown underneath. n=3; C. Representative Col I and Col III immunofluorescence images of lung tissue in the mice exposed to FPM for 7 days. Scale bar = 100 μm. Results are shown as mean ± SD. n.s. indicates no statistical significance.

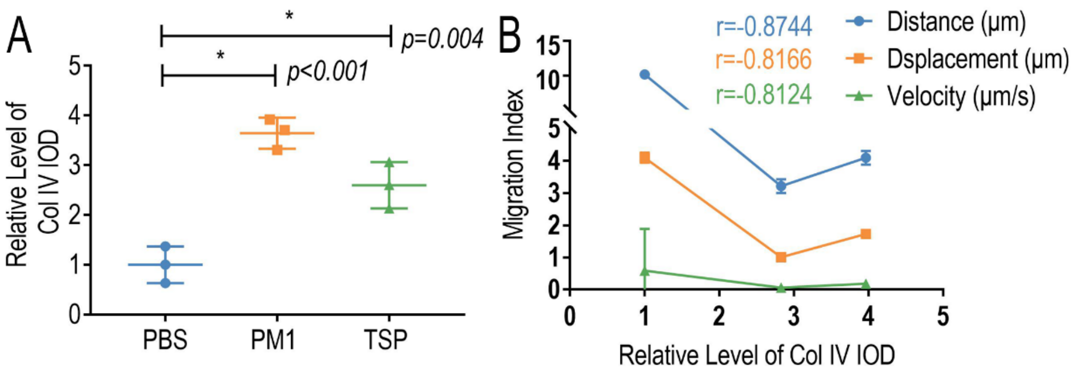

**Supplementary Figure S18. A.** The integrated optical density (IOD) of Col IV immunofluorescence images of lung tissue exposed to FPM for 7 days, based on Image J analysis. n=3. Results are shown as mean ± SD. *p<0.05 after ANOVA with Dunnett's tests. **B.** Pearson's correlation line of the migration index (migration distance, displacement and velocity) of different groups with integrated optical density (IOD) of Col IV in corresponding lung tissue slice as panel A. n=3. Results are shown as mean ± SD. *p<0.05 after ANOVA with Dunnett's tests.

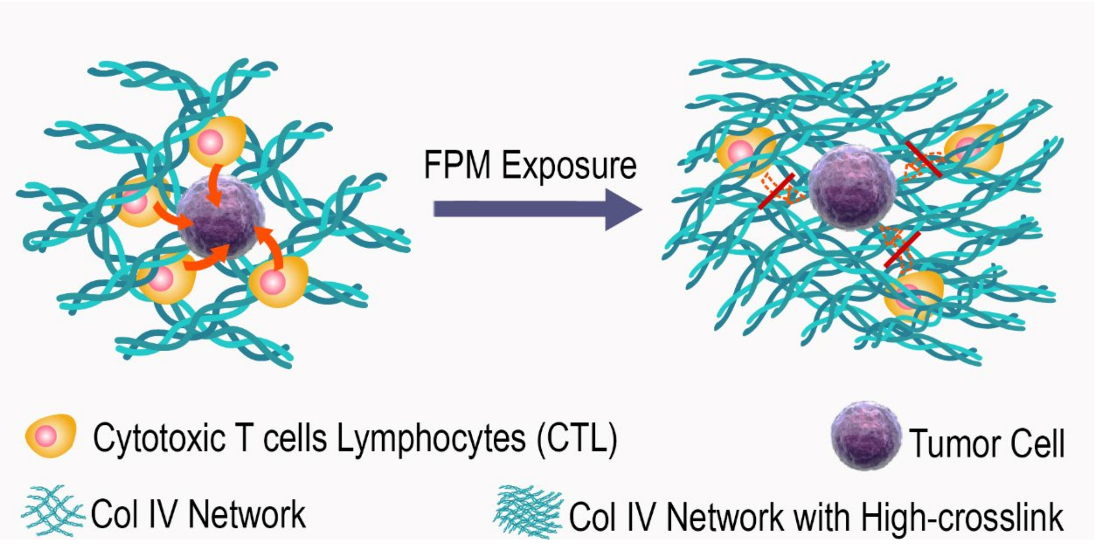

**Supplementary Figure S19.** Schematic diagram about CTLs' migration on the lung tissue exposed to FPM.

**Supplementary Video S4-7.** Dynamic migration videos of T cells on FPM-exposed lung tissue pre-treated with collagenase D (0.05 mg/mL) for 5 min. Jurkat T cells were pre-stained with Calcein-AM and labelled as green in videos. Collagen IV in lung tissue were labelled as red.

### 6. The Effect of FPM on Col IV Crosslink.

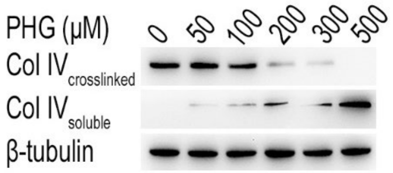

**Supplementary Figure S20.** Western blotting analysis of 'soluble' and 'crosslinked' Col IV fraction in M2-10B4 cells lysate after the cells were treated with different concentration of crosslink inhibitor phloroglucinol (PHG) for 7 days.

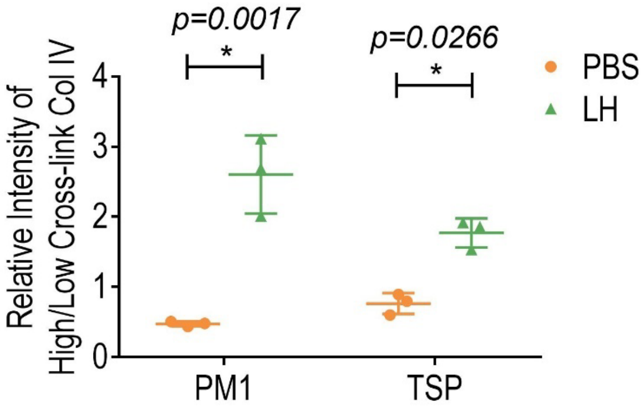

**Supplementary Figure S21.** The relative intensity of high-crosslink Col IV to low-crosslink ones according to the WB results in M2-10B4 cells lysate enriched with soluble collagen after their treatment with FPM or the mixture of lung homogenate (LH) and FPM, that is, LH-FPM. n=3. Results are shown as mean ± SD. *$p<0.05$ after ANOVA with Dunnett's tests.

### 7. Detection of Tryptic Peptides Containing Crosslink Site.

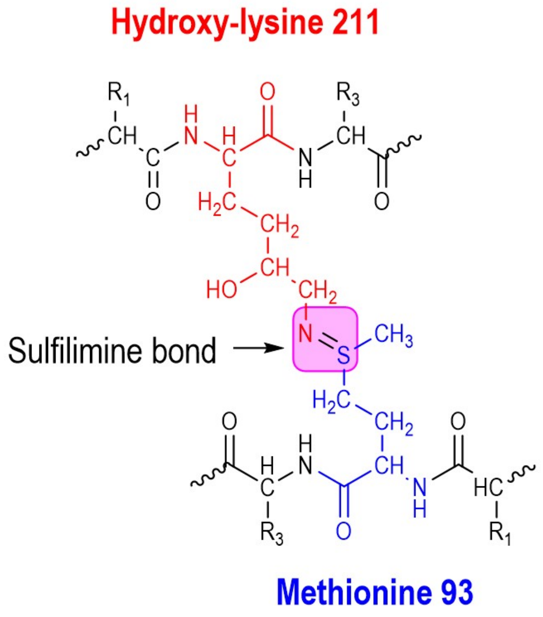

**Supplementary Figure S22.** Structure of sulfilimine bond formed at the covalent crosslinks of NC1 domains, shown in the lilac box.

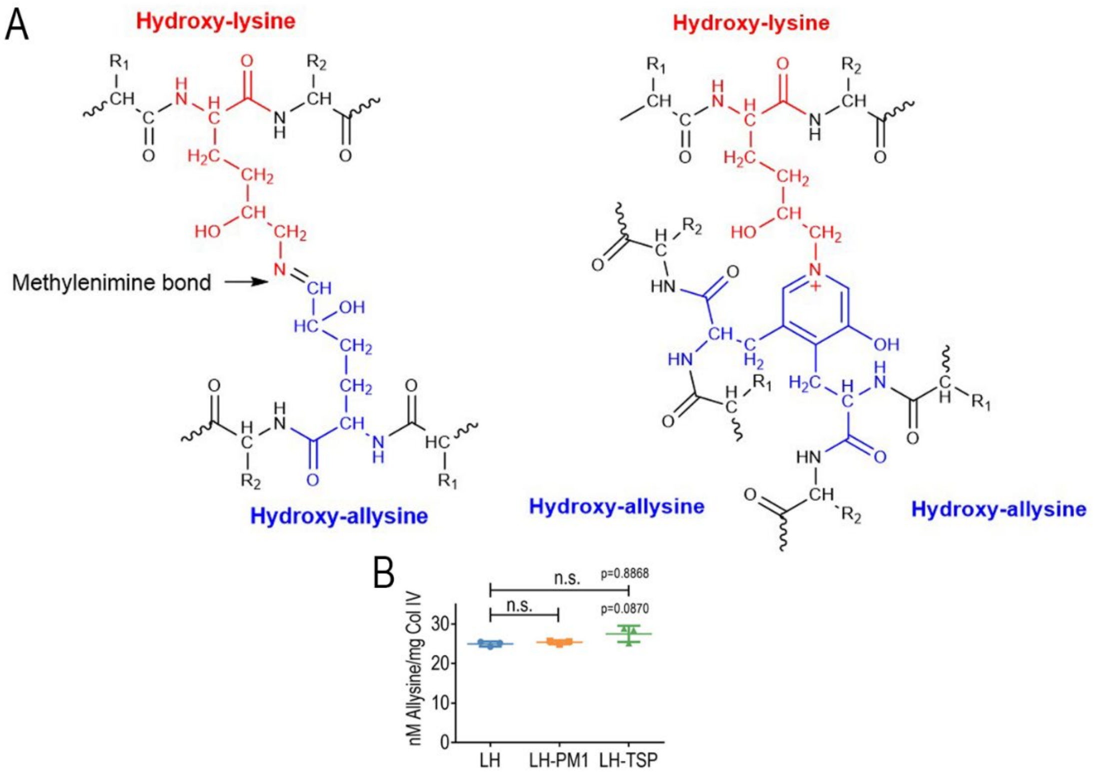

**Supplementary Figure S23.** A. The potential crosslink site formed at 7S domain, containing methylenimine bond (-C=N-, left) or pyridinum crosslink (right), shown in the lilac box; B. The allysine generated during the Col IV crosslinking after soluble Col IV were incubated with LH per se or the mixture of LH-FPM (LH-PM 1 and LH-TSP) for 30 min, detected with specific probes for allysine, with a serial content of oxidized bovine serum albumin containing known aldehydes as the internal standard (6). n=3. Results are shown as mean ± SD. n.s. indicates no statistical significance.

### 8. Preparation and Characterization of Protein Corona.

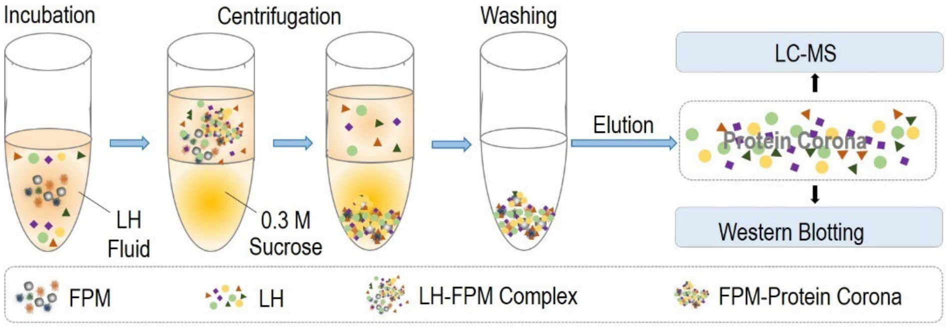

**Supplementary Figure S24.** Schematic diagram of separation and preparation of FPM' protein corona in lung homogenate (LH).

**Supplementary Table S2.** List of protein component identified by liquid chromatography-mass spectrometry (LC-MS) for PM1's and TSP's protein corona. Single Results Table respectively are given.

### 9. Effect of FPM on PXDN's Enzymatic Activity and Phase Separation.

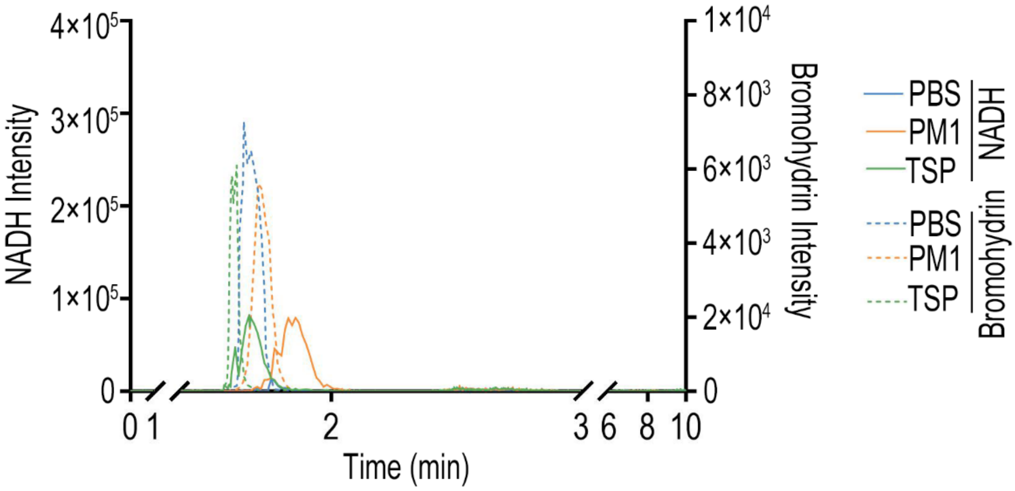

**Supplementary Figure S25.** Liquid chromatography-mass spectrometry (LC-MS) spectrum for NADH (dotted line) and the bromohydrin (line), according to the reported literature (1). The analysis was performed after the enzyme PXDN was incubated with FPM for 30 min and then catalyzed in the presence of 100 μM $H_2O_2$ and 200 μM NaBr at 37°C for 30 min.

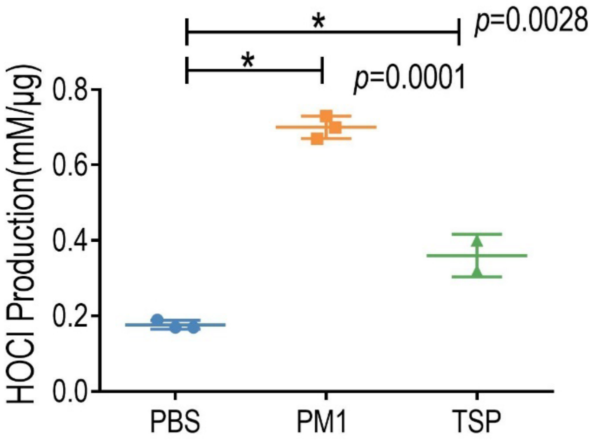

**Supplementary Figure S26.** HOCl production induced by PXDN measured with HClO detecting fluorescent probes, after the enzyme was incubated with FPM for 30 min and then catalyzed in the presence of 100 mM $H_2O_2$ and 200 mM NaCl, with a serial concentration of HClO as internal control. n=3. Results are shown as mean ± SD. *$p<0.05$ after ANOVA with Dunnett's tests.

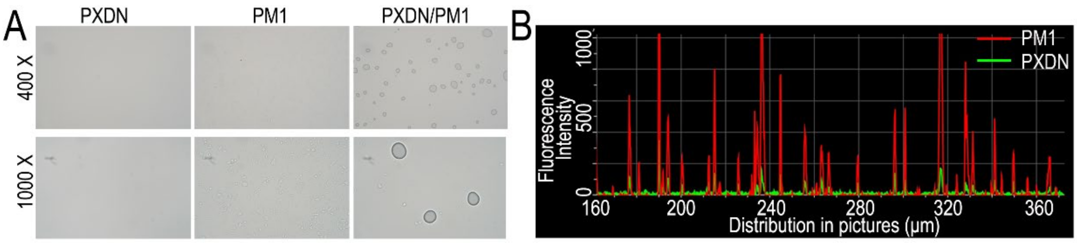

**Supplementary Figure S27. Effect of FPM on PXDN's phase separation.** A. Phase contrast microscopy of PXDN *per se* or incubated with FPM for 30 min in LH; B. The fluorescence distribution profiles of the cross-sectional region of liquid-like droplets on the FPM after FITC-labelled PXDN was incubated with rhodamine-labelled FPM in LH for 30 min.

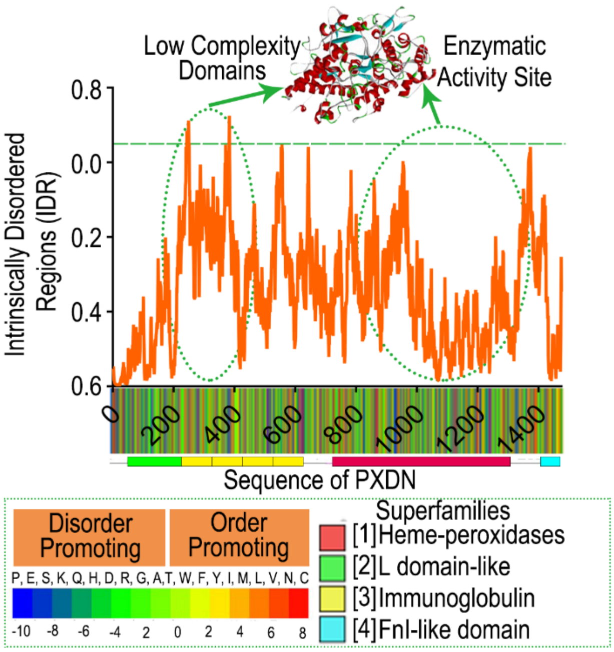

**Supplementary Figure S28.** Intrinsically disordered regions (IDRs) analysis of PXDN domains by the IUPred algorithm. The low complexity domains and enzymatic activity site were respectively labelled with dotted green circles. PXDN's template crystal structures were shown on the upper. Besides, bioinformatics analysis of the amino acid sequence of full-length PXDN shown underneath. The kinds of amino acids with order or disorder potential were listed with different color and the superfamily of PXDN was shown on the lower right (2).

### 10. Effect of PXDN Inhibitor on Lung Tissue Microenvironment and Lung Tumorigenesis.

#### 10.1 Effect of PXDN Inhibitor on Lung Tissue Structure.

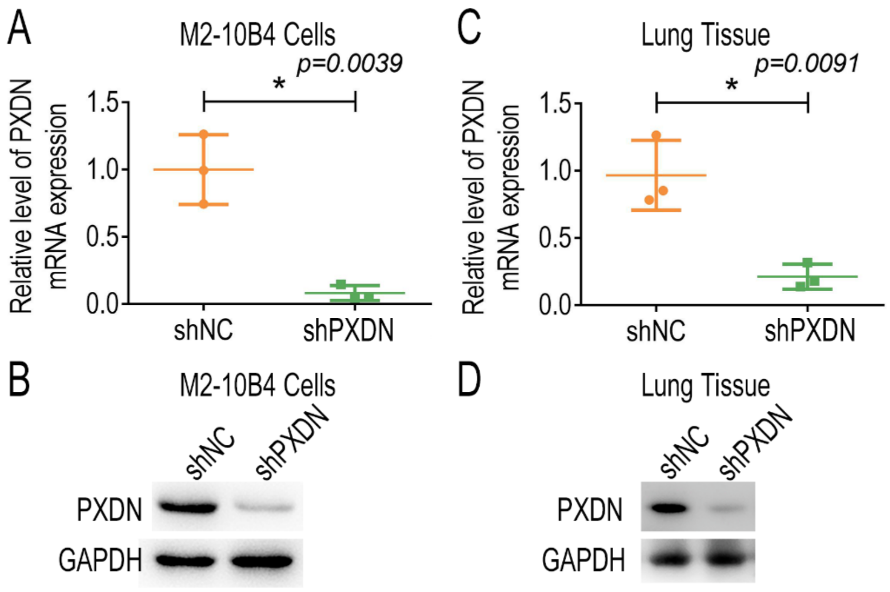

**Supplementary Figure S29. A and B.** The transcriptional level analysis (**A**) and Western blotting analysis (**B**) of PXDN in M2-10B4 cells after they were transfected with plasmids capable of ectopically expressing PXDN specific short hairpin RNA (shPXDN) or control shRNA (shNC) for 48 h; **C and D.** The transcriptional level analysis (**C**) and Western blotting analysis (**D**) of PXDN in lung tissue after the mice were administrated with 4 μg plasmids capable of ectopically expressing shPXDN mixed in the *in vivo*-jetPEI gene transfer regent through trachea injection every 3 day for 4 times.

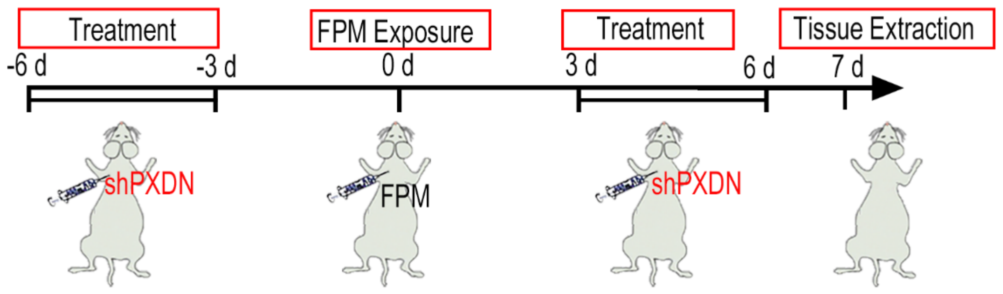

**Supplementary Figure S30.** Schematic diagram of analyzing the effect of PXDN specific short hairpin RNA interference (PXDN shRNA) on the structure of FPM-exposed lung tissue.

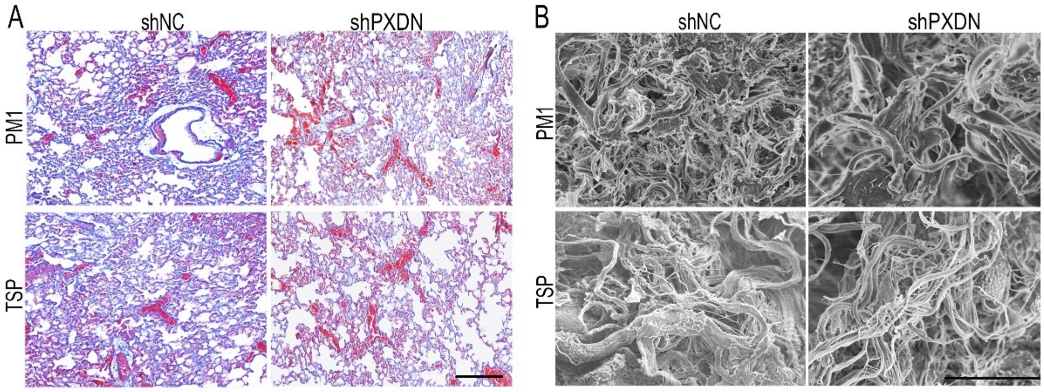

**Supplementary Figure S31. A.** Representative Massons' trichrome histological analysis of lung tissue in FPM-exposed mice administrated with shPXDN. Scale bar = 100 μm; **B.** Representative SEM images of interstitial matrix in the lung tissue in FPM-exposed mice administrated with PXDN shRNA. Scale bar = 50 μm.

### 10.2 Effect of PXDN Inhibitor on T Cell Migration.

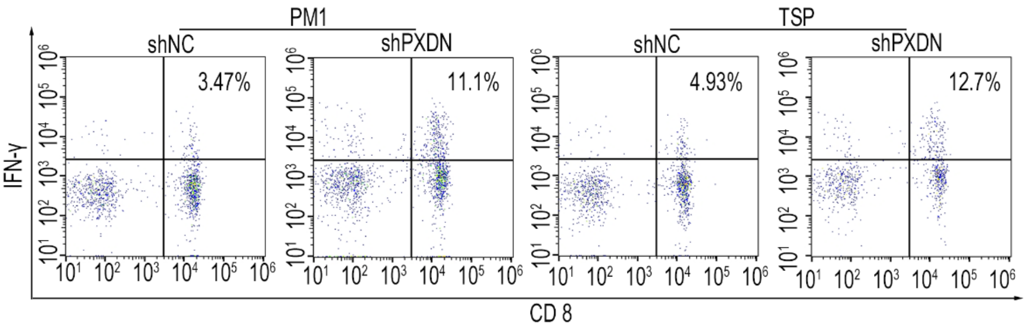

**Supplementary Figure S32.** Representative flow cytometry analysis of CTLs' infiltration (IFN-$\gamma^+$CD8$^+$/CD45$^+$CD3$^+$) into lung tissue of FPM-exposed group pretreated with shPXDN 1 day after they were stimulated with the LLC.

### 10.3 Effect of PXDN Inhibitor on Lung Tumorgenesis.

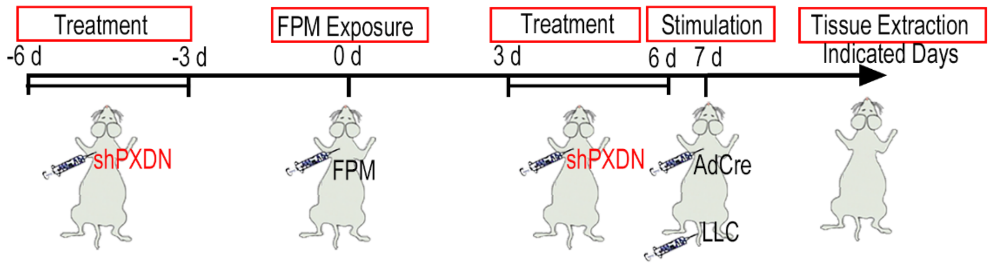

**Supplementary Figure S33. A.** Schematic diagram of analyzing the therapeutic effect of shPXDN on LLC-stimulated or Kras$^{G12D}$/p53$^{-/-}$-transgenic lung cancer model with FPM exposure.

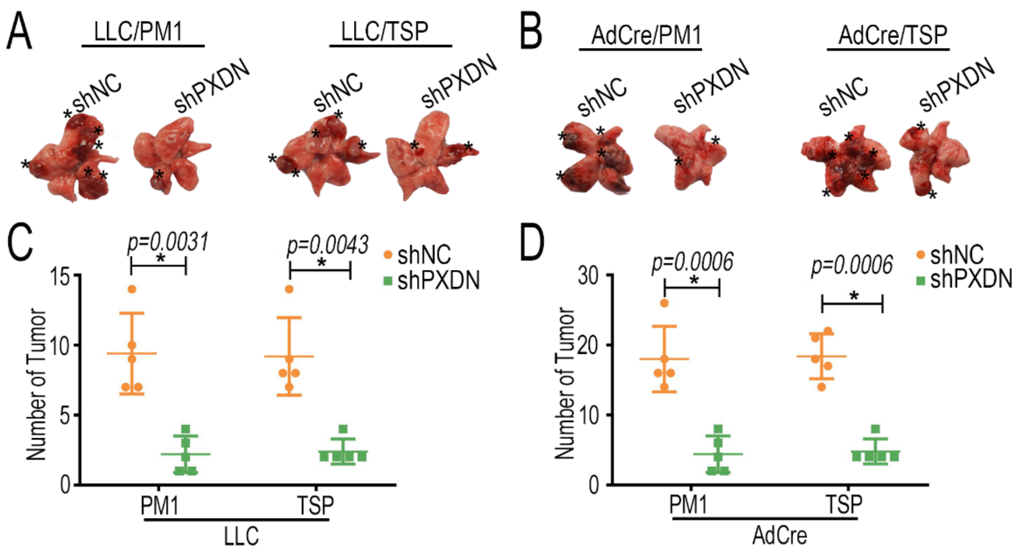

**Supplementary Figure S34. A and B.** Gross lung tissue images in LLC-stimulated (**A**) or Kras$^{G12D}$/p53$^{-/-}$-transgenic lung cancer model (**B**) administrated with shPXDN; **C and D.** Statistical analysis of tumor number in LLC-stimulated (**C**) or Kras$^{G12D}$/p53$^{-/-}$-transgenic lung cancer model (**D**) administrated with shPXDN. n=5. Results are shown as mean ± SD. **p*<0.05 after ANOVA with Dunnett's tests.

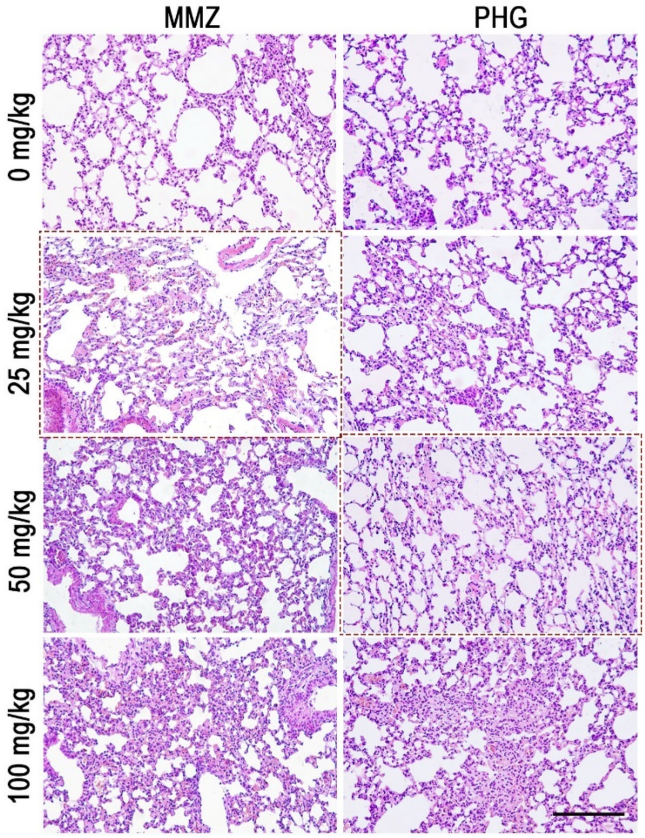

**Supplementary Figure S35.** Representative H&E staining images of lung tissue with intratracheal injection with different concentration of methimazole (MMZ) and phloroglucinol

(PHG) for 3 days. Scale bar = 100 μm. The concentration at the indicated group with red dotted lines was chosen as the subsequent administration dose.

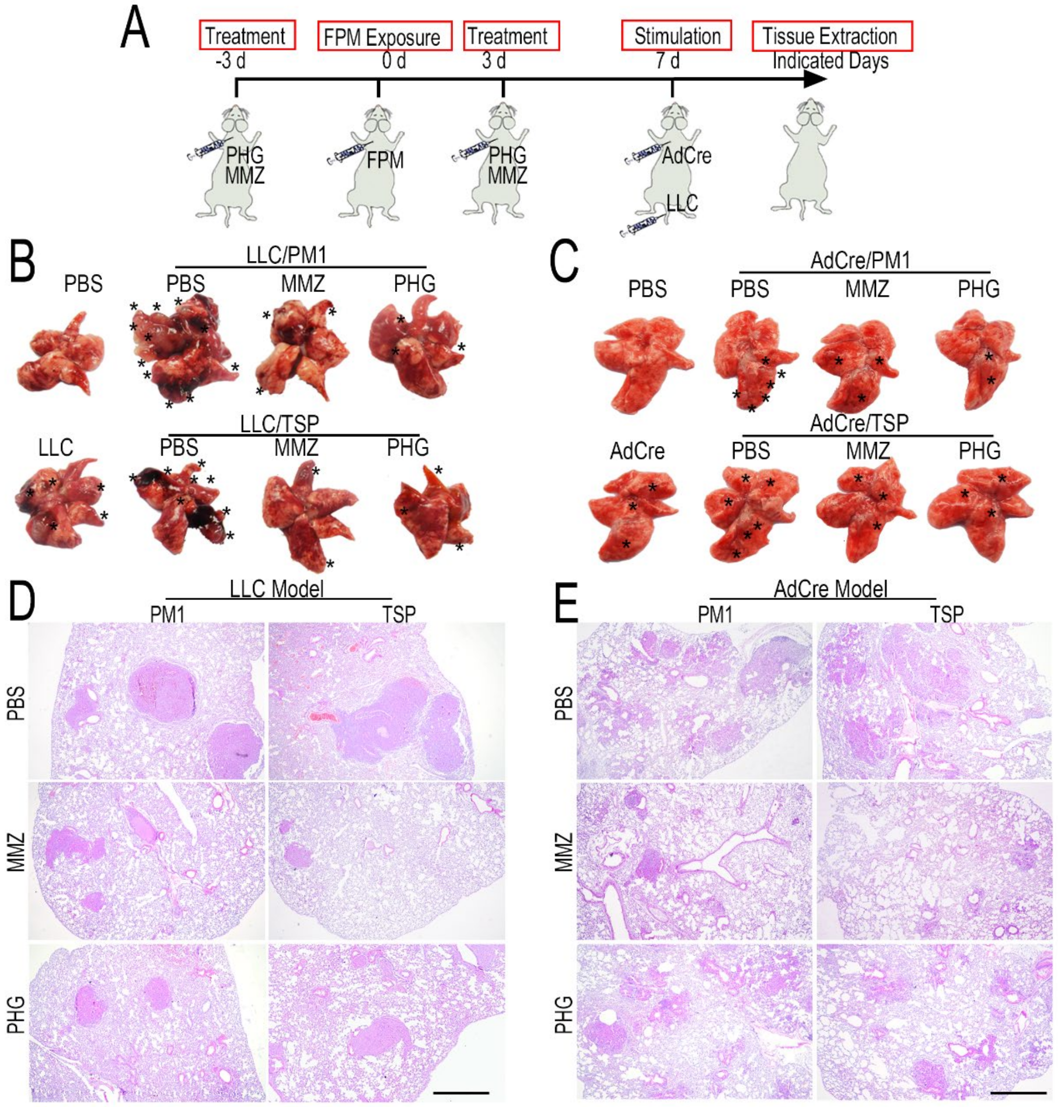

**Supplementary Figure S36. A.** Schematic diagram of analyzing the therapeutic effect of PXDN inhibition (MMZ or PHG) on LLC-stimulated or $Kras^{G12D}/p53^{-/-}$-transgenic lung cancer model with FPM exposure; **B and C.** Gross images of lung tissue yielded from LLC-induced model (**B**) or $Kras^{G12D}/p53^{-/-}$-transgenic lung cancer model (**C**) 20 days or 50 days after mice pretreated with MMZ or PHG were stimulated with LLC or AdCre; **D and E.** Representative H&E staining images of lung tissue yielded from LLC-induced model (**D**) or $Kras^{G12D}/p53^{-/-}$-transgenic lung cancer model (**E**) as panel B and C. Images are representative for three independent experiments.

### 11. Supplementary Methods

**Characterization of FPM.** A series of tests were performed to thoroughly characterize the nanoparticles. First, to analyze morphology of particles, dried PTFE filter membranes containing FPM was conducted with SEM microscope LEO1530VP (JEOL

Ltd., Tokyo, JAPAN). Second, all the particles were characterized for their Zeta potential and particle size using NanoSight NS300 instrument (Malvern Instruments, Malvern, UK). Third, to obtain essential information on the nanoparticles' size and shape, transmission electron microscopy (TEM) was carried out. After a few drops of deionized water-dispersed nanoparticles was dropped on the 300-mesh carbon-coated copper grid, TEM images of each sample were collected using TEM (JEOL Ltd.). Besides, the elemental analysis was performed on an element analysis instrument (Vario Micro Cube, Elementar, Germany), with the top 15 was listed.

**Cell Preparation and Culture.**

*Cell lines' culture*. Lewis lung cancer cell lines (LLC), mouse bone marrow fibroblasts M2-10B4 and Jurkat T cells were obtained by Stem Cell Bank, Chinese Academy of Sciences (Shanghai, China). LLC cells expressing ovalbumin peptide residues 257-264 ($OVA_{257-264}$) in the context of $H2K^b$ (OVA-LLC) were kindly provided by K. Zeng (Nanjing University, China). Cells were cultured in DMEM or RPMI 1640 medium containing 10% foetal bovine serum (Thermo Fisher Scientific, MA, USA), harvested at ~ 80% confluency, washed twice with phosphate buffer saline (PBS) and subcultured for passage. The short tandem repeat (STR) profiling of these cell lines was authenticated (Beijing Microread Genetics Co., Ltd, Beijing, China). And all cell lines were detected negative for mycoplasma contamination (Corues Biotechnology, Nanjing, China).

*Separation of primary cytotoxic $CD8^+$ T lymphocytes (CTLs)*. To exact the CTLs from lung tissue, the lung tissues in the FPM-exposed mice stimulated with LLC or OVA-LLC for indicated days, were respectively collected and digested with 2 mg/mL collagenase type I and IV (Thermo Fisher Scientific) for 20 min. Then a single-cell suspension was prepared using the program m_lung_02.01 on the gentleMACS™ Dissociator (Miltennyi Biotec, Bergisch Gladbach, Germany). the CTLs were isolated from this single-cell suspension using the $CD8^+$ T cell isolation kit with a MidiMACS™ separator (Miltennyi Biotec).

*Cloning, expression, and purification of peroxidasin (PXDN).* His-tagged full-length mouse peroxidasin homolog encoded on indicated vector was generated by GenScript and transfected into HEK 293 F cells using Lipo2000 (Invitrogen) according to standard selection and cultivation procedures with minor modifications (5). During large scale, cells were cultivated in Expi293™ Met (-) Expression Medium (Thermo Fisher Scientific). The harvested media were stored at 4 °C and eventually purified using Ni-NTA Agarose (Qiagen). Fractions with the best purity number were pooled, concentrated, and desalted using Centricon with a 100-kDa cutoff membrane (Millipore).

**Stimulation of FPM on LLC and Cytotoxic T Lymphocytes (CTLs).** To analyze the effect of FPM on the proliferation of LLC, cell counting kit-8 (CCK-8) test was performed. Briefly, $1\times10^{4}$ LLC were seeded in 96-well culture plates and then simulated with different concentration of FPM (0 μg/mL, 5 μg/mL, 10 μg/mL, 30 μg/mL, 50 μg/mL, 100 μg/mL and 500 μg/mL) and cultured for 24 h and 48 h. CCK-8 kit (DOJINDO LABORATORIES, Kumamoto, Japan) was used to examine the proliferation of LLC at indicated time points, with cells treated with 1×PBS as control. To further analyze the effect of FPM on the migration potential of T cells, the CTLs were stimulated with 10 μg/mL FPM, which showed slight cytotoxic, for 48 h.

**Extraction of Soluble Collagen IV.** To generate soluble collagen IV, the fused mouse bone marrow fibroblasts M2-10B4 cells were plated at high density and maintained at confluency for 7 days in the presence of 50 μg/mL ascorbic acid (Sangon Biotech, Shanghai, China), with media changes every 24 – 36 h. Crosslinking was inhibited by supplementing the culture conditions with indicated concentration (0, 50, 100, 200, 300 and 500 μM) of PHG. PHG and ascorbic acid treatments were initiated upon confluency. With the 200 μM PHG, which could be sufficient to inhibit the Col IV crosslink, after the M2-10B4 cells stimulated for 7 days and collected through scrape, cultured cells and matrix were homogenized in 1% (w/v) deoxycholate (Aladdin) with sonication, and the insoluble material isolated after centrifugation at 20,000 × *g* for 15 min. Then the pellet was lysed with RPMI (Beyotime Biotechnology, Shanghai, China) at ice for 30 min. The supernatant containing soluble Col IV was collected after centrifugation at 20,000 × *g* for 10 min and then incubated with 1 mg/mL FPM per se, 1 mL lung homogenate (LH) or the mixture of FPM-LH (with the volume ratio of 1:10) for 4 h.

Then the samples were collected for further Western boltting (WB) analysis to detect the change of Col IV crosslink. Besides, to perform cellular experimental analysis, the M2-10B4 cells were incubated with 200 μM PHG for 24 h and then treated with the same stimulation for 24 h. The cell samples were collected for further immunofluorescence (IF) analysis.

**Analysis of Crosslinking Extent of Different Collagen.** Collagen crosslink was assessed biochemically by separating different collagen fractionation via serial extractions, including neutral salt (freshly secreted collagens and procollagens), acetic acid (more mature collagens), and acid pepsin (fibrillar, moderately crosslinked collagens) and insoluble high crosslink ones from fresh lung tissue as literature reported (3; 4). Briefly, the whole lung tissue was homogenized in neutral salt buffer (0.5 M NaCl, 0.05 M Tris, pH 7.5; Sangon Biotech) and incubated at 4 °C overnight on a rotary shaker. After centrifugation at 24,000 × *g* for 30min, the supernatant was collected (fraction A: neutral salt-soluble collagen). The resulting pellet was then extracted with 0.5 M acetic acid (fraction B: acid-soluble collagen; Sangon Biotech), followed by pepsin (2 mg/mL in 0.5 M acetic acid, fraction C: pepsin-soluble collagen; Sangon Biotech). The remaining insoluble fraction D represents mature, highly crosslinked collagen. Then Type I, III and IV collagens with different extractions were analyzed by corresponding enzyme linked immunosorbent assay (ELISA) kits (Nanjing Jiancheng Bioengineering Institute, Nanjing, China). The level of collagen crosslink was calculated as collagen in fraction D divided by total collagen summed by faction A, B, C and D.

**Detection of Allysine at 7S domain Crosslinking Site.** For 7S domain, the detection of primary product allysine could reflect the level of its crosslink. With the reported specific and efficient probes to allysine (6), crosslink of the soluble Col IV incubated with LH or the mixture of LH-FPM was respectively analyzed. Briefly, the 5 mg/mL soluble collagen IV was incubated LH or LH-FPM mixture and 5 mM probes, for 30 min at 37 °C. Then the fluorescence intensity was detected with the exciting wavelengths at 488 nm on the microplate reader (Thermo Fisher Scientific). To be estimated, to quantify the yielded allysine, a serial content of oxidized bovine serum albumin (BSA) containing known aldehydes as the internal standard (oxidized BSA: 16 nM aldehyde/mg; BSA: 1.2 nM aldehyde/mg). For the oxidized BSA, sodium aspartate (13 mg) was added to 50 mg/mL BSA in phosphate buffered saline (PBS), followed by the addition of a solution of ferric chloride (10 μL, 10 mM) and left to stir at room temperature overnight. A BSA protein standard without the addition of ferric chloride was run in parallel as a control.

**Preparation and Separation of FPM's Protein Corona.** First, lung homogenate (LH) was extracted from the lungs of health mice according to institutional bioethics approval. Briefly, the extracted lung tissue samples were homogenized in equal volume of PBS by a homogenizer (approximately 3 mice/mL LH), and then centrifuged to remove the debris to obtain LH. 10 mg/mL FPM were incubated with LH at the volume ratio of 1:10 under stirring at 4 °C for indicated time. Then, the mixture was centrifuged through a 0.3 M sucrose cushion for 20 min at 4 °C at 15,300 × *g*, in order to separate the nanoparticle-corona complexes. Then, after rinsed with 1 × PBS for 3 times, proteins in the corona were eluted by adding RIPM lysis buffer (50 mM Tris pH7.4, 150 mM NaCl, 1% Triton X-100, 1% deoxycholate, 0.1% SDS) to the pellet on the ice for 1 h. After centrifugation (20 min at 15,300 × *g* at 4 °C), the supernatant enrich protein corona was collected and store at -20 °C.

**LC-MS Analysis and Database Searches of Protein Corona.** The protocol to analyze protein with LC-MS adhered to a method described previously (7; 8). Briefly, after samples were quantified with bicinchoninic acid protein assay (BCA) kit; 100 μg total protein was reduced by adding 1 M DL-Dithiothreitol (DTT) (Sigma-Aldrich) (60 °C, 1 h), and free cysteines were alkylated with 1M iodoacetamide (IAA) (Sigma-Aldrich) (room temperature, 10 min in the dark). The alkylated proteins were centrifuged in the 10K ultrafiltration tube (Thermo Fisher Scientific), and the proteins were retained in the 10K ultrafiltration tube. The proteins were further washed with 100 mM tetratehylammonium bromide (TEAB) for three times at 4 °C for 20 min by centrifugation at 12,000 rpm. Then the protein was digested with 2 μg porcine sequencing grade trypsin (LC-MS Grade, Sigma-Aldrich) overnight at 37 °C. After digestion, the resulted peptides were collected (12,000 rpm, 20 min, 4 °C), desalted by

Zeba Spin Desalting Columns (Thermo Fisher Scientific) and further enriched by C18 reversed-phase columns (Epoch Life Science, Missouri City, Texas, USA). The samples were then subjected to LC-MS analysis. To identify the composition of protein corona, identification of peptides and proteins from continuum LC-MS data was performed with the ProteinPilot™ 4.5 software (AB SCIEX). Proteins were analyzed by searching the mouse taxon of the UniProtKB/SwissProt database (release 2011_11). The proteins with at least one specific high-scoring peptides were detected and exported from ProteinPilot for the final LC-MS data file at the protein level.

**Flow Cytometry Analysis.** Lung tissue were digested with 2 mg/mL collagenase type I and IV (Thermo Fisher Scientific) for 30 min to generate a single-cell suspension. Cell suspensions were filtered through 70 μm cell strainers, and red blood cells were lysed. $1\times10^6$ cells/ml were treated with cell activation cocktail (BioLegend, San Diego, California, USA) for the intracellular staining according to the manufacturer's protocol. After cells were washed with PBS containing 1% BSA, cells were blocked with 1% BSA at 4 °C for 30 min. Zombie Violet™ Fixable Viability Kit was used for live/dead cell determination. Then cells were stained on ice for 30 min with surface-staining antibodies, FITC anti-mouse CD45, BV711 anti-mouse CD3, APC anti-mouse CD8a, and then washed, fixated and permeabilised with the fixation/permeabilization solution kiT (BD Biosciences, San Jose, CA, USA) and stained with cytokine PE anti-mouse interferon gamma (IFN-γ) antibodies. in the dark for 30 min at 4 °C. The samples were centrifuged at 400 – 500 × *g* for 5 min at 4 °C to remove unbound antibody. After rinsing for 3 times, each sample was re-suspended for analysis using a BD Fluorescence Activated Cell Sorter (FACS) Calibur (BD Biosciences). Unconjugated antibodies and IgG controls were run in parallel to set the background. All antibodies and their isotype control antibodies were obtained from BioLegend (San Diego, CA, USA).

**Western Blotting.** According to the standard protocol, different proteins were separated by SDS-PAGE. To be estimated, the proteins in corona from the nanoparticles were eluted with equal and adequate PAGE sample buffer containing 1 mM phenylmethanesulfonyl fluoride (PMSF) (Sigma-Aldrich) and same volume of eluted corona proteins was analyzed. Besides, to further estimate the content of PXDN adsorbed on the FPM, different amount of PXDN (100 ng, 500 ng, 1,000 ng and 2,000 ng) together with the corona protein samples was separated by SDS-PAGE and analyzed by western blotting. Then the proteins were transferred onto the polyvinylidine difluoride (PVDF) membranes (Bio-Rad, California, USA). The membranes were blocked with skim milk and then incubated with primary antibody – PXDN (Merck Millipore), Type I collagen (Col I, Boster Biological Technology co.ltd, Wuhan, China), Type III collagen (Col III, ABclonal Technology, Wuhan, China), Type IV collagen (Col IV, Abcam, Cambridge, MA) and glyceraldehyde-3-phosphate dehydrogenase (GAPDH, Abcam) at 4 $^{\circ}$C with gentle shaking overnight. After washed with PBST (PBS with 0.1% Tween-20) for 5 times, the membrane was incubated with horseradish peroxidase-conjugated anti-rabbit, anti-mouse or anti-goat IgG (Life Technologies, Grand Island, NY, USA) at room temperature. After rinse, positive signal was visualized using an enhanced chemiluminescence system (Cell Signaling Technology). The band intensity was quantitated using image J software (http://rsb.info.nih.gov/ij/) and the statistical analysis of three independent experiments was performed.

**RNA Isolation and Quantitative Real-time PCR.** RNA of cells or lung tissues were extracted by using Trizol reagent (Life Technologies). For mRNA detection, real-time polymerase chain reaction (PCR) was launched in an ABI 7300 Fast Real-time PCR System (Applied Biosystems, FosterCity, CA) using the SYBR Prime Script RT-PCR Kit (Takara Bio, Shiga, Japan). Each sample was analyzed in triplicates and repeated for three or four independent assays with β-actin as internal control. Primers of integrin-1 (ITGB1), C-X-C motif chemokine receptor 3 (CXCR 3), Rho-associated kinase (ROCKi) and peroxidasin (PXDN)are listed as follows (Shanghai Generay Biothech Co., Ltd, Shanghai, China):

ITGB1-Forward: 5'- CGTGGTTGCCGGAATTGTTC -3',

ITGB1-Reverse: 5'- ACCAGCTTTACGTCCATAGTTTG -3;

CXCR3-Forward: 5'- TACCTTGAGGTTAGTGAACGTCA -3',

CXCR3-Reverse: 5'- CGCTCTCGTTTTCCCCATAATC -3';

ROCKi-Forward: 5'- AACATGCTGCTGGATAAATCTGG -3',

ROCKi-Reverse: 5'- TGTATCACATCGTACCATGCCT -3';

PXDN-Forward: 5'- CCTGTGTTTCCGTACCACCG -3',

PXDN-Reverse: 5'- CTCTGATTCTGTTGAACCGAAGA -3';

β-actin-Forward: 5'- GGCTGTATTCCCCTCCATCG-3',

β-actin- Reverse:5'- CCAGTTGGTAACAATGCCATGT-3'.

**Histological Studies.** The lung tissue fixed in 2.5% paraformaldehyde (PFA) was embedded in paraffin and cut into sections for the hematoxylin and eosin (H&E) and Masson's trichrome (Abcam) staining (NanJing KeyGen Biotech Co.,Ltd., Nanjing, China) according to the manufacturer's instructions with slight modification. Stained sections were photographed at different magnification under a microscope. Under blindfold conditions with standard light microscopy, tumor burden (based on the percentage of the area of tumor regions *versus* that of total lung) according to H&E-stained sections of all five lung lobes was quantified with ImageJ software. Besides, to observe the interstitial ECM structure, lung tissues were fixed with glutaraldehyde at 4 °C for 48 h, dehydrated with an ethanol gradient and dried at the critical point. Then the samples were sprayed with gold particles and observed with SEM (SFEG Leo 1550, AMO GmbH, Aachen, Germany).

**Immunofluorescence Staining.** Lung tissue samples were collected, frozen at optimal cutting temperature (OCT) medium (Thermo Fisher Scientific) and cut into sections. The sections or M2-10B4 cells incubated with 10 μg/mL rhodamine-labelled FPM and 1 μg/mL PXDN (at the volume ratio of 1:10) for 1 h were fixed with 4% paraformaldehyde (PFA, Sigma-Aldrich) and stained with primary antibody at 4 °C overnight. The primary antibodies, including PXDN, Col I, Col III, Col IV and CD8. Next, the sections were incubated with secondary antibody Alexa Fluor (Life Technologies) for 1 h at room temperature, followed by 4,6-diamidino-2-phenylindole (DAPI, Beyotime) for nuclear staining. And then the sections were imaged by LSM

980 with Airyscan 2 confocal microscope (Carl Zeiss; Oberkochen; Germany). To further characterize the crosslink level, based on IF mages of Col IV, look-up tables (LUTs) analysis based on the fluorescence intensity, surface plot analysis based on the invert binary distance of fluorescence distribution, were respectively accomplished employing Image J (Image J Software, National Institutes of Health, Bethesda, MD, USA). Using the 'ridge detection' plugin in Image J, binary images of Col IV network was generated, and the related quantitative analysis of junction number and junction density were created and compared. Besides, the EdU positive percentage was analyzed by the Bioapps Tools in ZEISS ZEN 3.4 (Carl Zeiss).

**Enzyme-linked Immunospot Assay (ELISPOT).** The lung tissue exposed to PBS or FPM were excised after the intravenous stimulation of LLC-OVA cells for 1 day. After CTLs were separated, $1x10^5$ CTLs were added in each well of IFN-γ antibody pre-coated plate and stimulated by $4x10^4$ irradiated LLC-OVA cells or PMA for 24h in RPMI-1640 supplemented with 10% fetal bovine serum (FBS), 100 U/ml penicillin and 0.1 mg/ml streptomycin. IFN-γ producing CTLs were enumerated by a mouse IFN-γ precoated ELISPOT kit (Dakewe Biotech Co., Ltd.) according to the manufacture instructions. The results were analyzed by AID iSpot (AID-Autoimmun Diagnostika GmbH, Strassberg, Germany).
